# Evaluating the effectiveness of artifact correction and rejection in event-related potential research

**DOI:** 10.1101/2023.09.16.558075

**Authors:** Guanghui Zhang, David R. Garrett, Aaron M. Simmons, John E. Kiat, Steven J. Luck

## Abstract

Eyeblinks and other large artifacts can create two major problems in event-related potential (ERP) research, namely confounds and increased noise. Here, we developed a method for assessing the effectiveness of artifact correction and rejection methods at minimizing these two problems. We then used this method to assess a common artifact minimization approach, in which independent component analysis (ICA) is used to correct ocular artifacts, and artifact rejection is used to reject trials with extreme values resulting from other sources (e.g., movement artifacts). This approach was applied to data from five common ERP components (P3b, N400, N170, mismatch negativity, and error-related negativity). Four common scoring methods (mean amplitude, peak amplitude, peak latency, and 50% area latency) were examined for each component. We found that eyeblinks differed systematically across experimental conditions for several of the components. We also found that artifact correction was reasonably effective at minimizing these confounds, although it did not usually eliminate them completely. In addition, we found that the rejection of trials with extreme voltage values was effective at reducing noise, with the benefits of eliminating these trials outweighing the reduced number of trials available for averaging. For researchers who are analyzing similar ERP components and participant populations, this combination of artifact correction and rejection approaches should minimize artifact-related confounds and lead to improved data quality. Researchers who are analyzing other components or participant populations can use the method developed in this study to determine which artifact minimization approaches are effective in their data.

## 1. Introduction

Eyeblinks generate large artifacts in electroencephalographic (EEG) recordings, typically exceeding 200 µV at the frontal pole (Fp1 and Fp2) and 30 µV at the vertex (Cz; see Lins et al., 1993 for normative data). Large artifacts can also occur anywhere on the scalp as a result of movements, skin potentials, muscle contractions, and idiosyncratic events (see Luck, 2014 for an overview of common artifacts in ERP experiments). Consequently, almost all event-related potential (ERP) studies employ an artifact rejection and/or artifact correction approach to deal with these artifacts, and several guidelines for publishing ERP and time-frequency studies indicate that this is essential (Duncan et al., 2009; Keil et al., 2014, 2022; Picton et al., 2000).

The present study examines the effectiveness of a common approach to dealing with eyeblinks and other artifacts that produce extreme values, in which artifacts with stable scalp distributions are corrected using independent component analysis (ICA; Chaumon et al., 2015; Jung, Makeig, Humphries, et al., 2000; Jung, Makeig, Westerfield, et al., 2000) and any trials that have extreme voltage deflections in any channel in the corrected data are also excluded from the averaged ERPs (artifact rejection; Islam et al., 2016; Nolan et al., 2010). This general approach is used very widely: in the first 10 issues of the 2023 volume of *Psychophysiology*, there were at least 18 papers that used some variant of this general approach (see Addante et al., 2023; Arnau et al., 2023; Bruchmann et al., 2023; Chen & Chen, 2023; Fan et al., 2023; Hubbard et al., 2023; Lin et al., 2023; Liu et al., 2023; Morales et al., 2023; Nguyen et al., 2023; Nicolaisen-Sobesky et al., 2023; Paraskevoudi & SanMiguel, 2023; Ringer et al., 2023; Schmuck et al., 2023; Sun et al., 2023; Tao et al., 2023; Wood et al., 2023; Zheng et al., 2023).

A careful assessment of this combined correction-rejection approach is particularly relevant at this moment because the usefulness of extensive EEG preprocessing has been questioned (Delorme, 2023) and because a new metric of data quality is now available that is directly related to effect sizes and statistical power (Luck et al., 2021). The rejection of trials containing artifacts may be particularly problematic because it will decrease the number of trials included in the averages, making the averaged ERP waveforms noisier. The new metric of data quality takes into account both the single trial noise level and the number of trials being averaged together, making it possible to determine whether the benefit of eliminating noisy trials outweighs the cost of having fewer trials.

The goal of the present investigation was to assess the effectiveness of an artifact minimization approach that is widely used and easy to implement using both open-source and commercial analysis packages. If this approach works well, researchers can keep using it, avoiding the considerable work required to implement more complex approaches that might have only marginal benefits. If it works poorly, however, this would establish a need for developing and implementing better approaches. We did not attempt to answer the question of which approach to artifact minimization is best, which would be difficult given the sheer number of available correction and rejection approaches. We did, however, aim to provide a well-justified and straightforward method for assessing the effectiveness of different artifact minimization approaches. Researchers could apply this method to their own data to determine whether their current artifact minimization approach is effective (because, as we will show, the effectiveness of an approach depends on the nature of the data). Methodologists could use this method to assess the effectiveness of new or improved artifact minimization approaches.

### 1.1. Goals of Artifact Correction and Rejection

To evaluate an approach to rejecting or correcting artifacts, it is essential to begin by carefully defining the goals of rejection and correction. Previous examinations of ICA-based artifact correction with real data have focused primarily on theoretical goals, such as minimizing mutual information and obtaining independent components with dipolar scalp maps (Delorme et al., 2012; Hoffmann & Falkenstein, 2008; Klug & Gramann, 2021; Winkler et al., 2015). Here, we consider the narrower goal of assessing whether two or more conditions or groups truly differ from each other in the amplitude or latency of a specific ERP component. There are two main ways in which artifacts can interfere with this goal (Luck, 2014, 2022). First, and most importantly, artifacts can be a potential *confound*. For example, if participants blink more in one condition than in another, the electrooculographic (EOG) voltage produced by the blinks may create a difference in the ERP waveforms between the conditions. This difference may then be interpreted as a difference in EEG activity between conditions rather than as a difference in EOG activity, leading to an incorrect scientific conclusion.

Second, artifacts may be a source of *uncontrolled variance* that decrease statistical power and cause a true effect not to be statistically significant. For example, a few random trials with a ±200 µV movement artifact in each participant’s data could be enough to add substantial error variance and prevent a true effect from being statistically significant. This would also lead to an incorrect scientific conclusion (or, more precisely, a failure to provide sufficient evidence for the correct conclusion). The present study addresses both of these issues, examining whether artifact correction and rejection help to minimize consistent but artifactual differences between conditions (i.e., reduce potential confounds) and help to minimize uncontrolled variance (i.e., increase statistical power). Note that although some prior studies of the effectiveness of artifact correction have assessed uncontrolled variance or statistical power(e.g., Delorme, 2023; Klug & Gramann, 2021; Mennes et al., 2010), previous research has largely neglected the possibility that artifacts are a confound. It is unlikely that any artifact correction method will be perfect, and it is essential to assess whether any residual artifactual signals create meaningful confounds in a given study.

EOG signals are also sometimes used for a third purpose in studies with visual stimuli, namely ensuring that the eyes are open and pointed in the appropriate direction when the stimulus is presented (Luck, 2014, 2022). For example, even when ICA is used to correct for EOG artifacts resulting from blinks, it can still be useful to reject trials on which the eyes are closed at the time of the stimulus. This issue arises in a fairly large number of studies, so we will consider it in the present analyses. In addition, when lateralized stimuli are used, EOG signals can be used to demonstrate that the eyes did not deviate systematically from fixation and change the sensory input. This is a less common issue that requires special analytic approaches(Luck, 2022; Woodman & Luck, 2003), so we will not consider it in the present analyses.

### 1.2. The Present Study

Most previous studies of the effectiveness of artifact minimization approaches have assessed data from a single study, often with a relatively small number of participants. This makes it difficult to know whether the results would generalize to other studies. In the present study, we evaluated the effectiveness of artifact correction and rejection across a broad range of experimental paradigms with a reasonably large sample size. Specifically, we used the publicly available ERP CORE (Compendium of Open Resources and Experiments; Kappenman et al., 2021), which includes data from 40 young adults who performed six standardized paradigms that yielded seven commonly-studied ERP components: P3b, N400, N170, N2pc, mismatch negativity (MMN), error-related negativity (ERN), and lateralized readiness potential (LRP). We did not analyze the N2pc and LRP data, because ocular artifacts create very different issues for these components than for most ERP components^1^. For the other five components, we asked (a) whether eyeblinks differed across experimental conditions and were therefore a potential confound, (b) whether ICA effectively minimized this confound, (c) whether ICA decreased or increased the data quality, and (d) whether the rejection of trials with extreme values increased the data quality even though it reduced the number of trials included in the averaged ERP waveforms. We also examined the effects of rejecting trials with blinks that interfered with the ability to see the stimuli.

We did not examine the effectiveness of rejecting trials in which a blink occurred at any time in the epoch, which was the standard approach prior to the widespread adoption of blink correction methods. When that approach was standard, it was also typical to ask participants to minimize blinking or to blink during the intertrial interval. In the ERP CORE tasks, by contrast, participants were instructed that they could blink immediately after responding, which was typically within the epoch. This is not a situation in which researchers would typically reject trials with a blink at any point in the epoch, so examining the effectiveness of this type of artifact rejection approach using the ERP CORE data would not be a fair test of the approach.

Blink-related confounds are a potential problem in a large proportion of ERP studies, so they are the focus of the present investigation. Our method can also be directly applied to vertical eye movements, which have very similar scalp distributions to blinks. Other kinds of confounds may be of concern in other experimental paradigms or participant populations, and our method could be modified for those other confounds.

By examining data from five different ERP components, the present investigation could draw relatively general conclusions about the effectiveness of the common approach of combining ICA-based correction for blinks with artifact rejection to eliminate trials with extreme voltage deflections. Our results may not generalize to all experimental paradigms and participant populations, but they will still be of value for a relatively large number of ERP studies. Another major goal of the present study was to develop and assess a rigorous yet easy-to-implement method for assessing the effectiveness of artifact correction and rejection. This method could be utilized in future research to assess the effectiveness of alternative artifact minimization approaches and could be applied to other experimental paradigms and participant populations. Moreover, we sought to test the proposal that artifact correction and rejection have little or no value (Delorme, 2023).

Note that the present study focuses on conventional averaged ERPs. However, it would be straightforward to adapt our assessment method to other EEG analysis approaches, such as time-frequency analysis and multivariate pattern analysis.

### 1.3. Quantifying Systematic Confounds

In the ERP CORE data, blinks are the main artifacts that are likely to differ systematically across experimental conditions and create a confound. We assessed the possibility that blinks were a confound by examining the bipolar vertical electrooculogram (VEOG) signal prior to artifact correction. The eyeblink artifact appears to be primarily caused by the eyelid sliding across the cornea^2^ (Lins et al., 1993), which creates a large positive deflection above the eyes and a smaller negative deflection below the eyes. We computed the bipolar VEOG as the voltage above the eyes (which is positive during a blink) minus the voltage below the eyes (which is negative during a blink). A positive value minus a negative value creates a large positive value, so this bipolar derivation effectively magnifies the blink-related activity. In addition, most EEG activity is similar at electrodes under and over the eyes, so this difference subtracts out most (but not all) neural signals. Thus, the bipolar VEOG channel provides a large and relatively pure index of blink activity.

We used this channel to determine the proportion of trials that contained blinks in each of the two conditions that were used to define a given component (e.g., face trials and car trials for the N170, error trials and correct trials for the ERN). If the proportion differs across conditions, then the blinks are a potential confound. Even if the number of blinks is equal across groups or conditions, the timing of the blinks might differ, and this could also be a significant confound. For example, if participants blink earlier on correct trials than on error trials in an ERN experiment, this will create an early negative voltage followed by a later positive voltage at frontal electrode sites in an error-minus-correct difference waveform. This pattern could be mistaken for an ERN followed by an error positivity (Pe). Indeed, this is exactly what we observed in our flankers paradigm. We assessed differences in the timing of blinks by comparing the averaged bipolar VEOG waveforms across conditions.

To determine the effectiveness of ICA-based artifact correction at minimizing blink-related confounds, we reconstructed the bipolar VEOG signal after artifact correction. If the corrected bipolar VEOG signals are nearly identical across conditions, then it is reasonable to conclude that the blink correction was effective in minimizing blink-related confounds. By contrast, a significant difference in bipolar VEOG between conditions after correction suggests that the correction was not completely successful. However, even if the blink-related activity has been perfectly eliminated, there could be differences between conditions in the bipolar VEOG signal as a result of volume-conducted ERP activity. We used semipartial correlations to determine whether any residual activity in the corrected bipolar VEOG signal reflected a failure of artifact correction or instead reflected volume-conducted voltages from the ERP component of interest.

### 1.4. Quantifying Data Quality

To determine whether artifact correction and rejection improved the data quality, we used a newly developed metric of data quality called the *standardized measurement error* (SME; Luck et al., 2021; Zhang & Luck, 2023). The SME estimates the standard error of measurement for a given amplitude or latency score from an averaged ERP waveform. It takes into account the fact that a given type of noise will have different effects on data quality depending on the method used to score a component’s amplitude or latency. For example, peak amplitude scores are much more distorted by high-frequency noise than are mean amplitude measures. In addition, when baseline correction is applied, low-frequency noise will have a larger effect on amplitudes at long latencies than at short latencies. The SME therefore estimates the measurement error for a particular score. A separate SME value is obtained for each participant, and the values can then be aggregated across participants by taking the root mean square of the individual-participant values (the *RMS(SME)*). The resulting RMS(SME) is directly related to the effect size for a difference between conditions or groups, which in turn is directly related to statistical power (see Luck et al., 2021, for details). Finally, because SME values represent the measurement error for amplitude or latency scores obtained from averaged ERP waveforms, they naturally reflect the combined effects of the single-trial noise level and the number of trials being averaged together. Thus, the SME provides an excellent means of determining whether the benefit of rejecting trials with extreme values (i.e., eliminating large artifactual voltages) outweighs the reduction in the number of trials being averaged together.

## 2. Method

The data analysis procedures were implemented with MATLAB 2021b (MathWorks Inc), using EEGLAB Toolbox v2023.0 (Delorme & Makeig, 2004) combined with ERPLAB Toolbox v9.20 (Lopez-Calderon & Luck, 2014). The data and scripts are available at https://osf.io/vpb79/. The present analyses can also be conducted without scripting by using the EEGLAB and ERPLAB graphical user interfaces. Thus, it should be straightforward for other investigators to apply our method to their own data.

### 2.1. ERP CORE Data and Preprocessing

We assessed the effects of artifact correction and artifact rejection on ERP data quality using the ERP CORE dataset (Kappenman et al., 2021), which can be found at https://doi.org/10.18115/D5JW4R. Comprehensive information concerning the participants, paradigms, recording techniques, and analysis protocols can be obtained from the original paper. The following section provides a concise summary of the participants, recording techniques, and preprocessing procedures. Additionally, each specific ERP paradigm is briefly described in its corresponding section within the Results section.

The ERP CORE dataset contains data from 40 neurotypical college students (25 women, 15 men), who were recruited from the University of California, Davis community. All 40 participants were included in the present analyses, regardless of the number of artifacts or behavioral errors present. The only exception is that one participant was excluded from the ERN analyses because this participant had only two usable error trials.

The EEG was recorded using a Biosemi ActiveTwo recording system (Biosemi B.V., Amsterdam) with DC coupling, active electrodes, an antialiasing filter (fifth-order sinc filter with a half-power cutoff at 204.8 Hz), and a sampling rate of 1024 Hz. Single-ended signals were recorded from 30 scalp sites (FP1, F3, F7, FC3, C3, C5, P3, P7, P9, PO7, PO3, O1, Oz, Pz, CPz, FP2, Fz, F4, F8, FC4, FCz, Cz, C4, C6, P4, P8, P10, PO8, PO4, 02) along with horizontal and vertical electrooculogram electrodes (HEOG left, HEOG right, VEOG lower).

Several preprocessing steps had already been carried out on the data within the ERP CORE resource. Stimulus event codes were adjusted to account for the intrinsic delay of the video monitor, and the data were resampled at 256 Hz using an antialiasing filter set at 115 Hz. Following this, the data were referenced to the average of P9 and P10 electrodes, which are electrically similar to the left and right mastoids but tend to be more stable. The one exception was the N170 paradigm, where Cz was used as the reference prior to averaging, and then the data were re-referenced to the average of all scalp sites after averaging. For all components, we also created a bipolar horizontal electrooculogram (HEOG-bipolar) channel as HEOG-left minus HEOG-right, and we created a bipolar vertical electrooculogram (VEOG-bipolar) as FP2 minus VEOG-lower^3^. In addition, a noncausal Butterworth high-pass filter with a half-amplitude cutoff at 0.1 Hz and a low-pass filter at 30 Hz (a roll-off of 12 dB/octave for both filters) were applied to the continuous EEG data.

Note that in all of the ERP CORE paradigms except the MMN task, participants were instructed to avoid blinking between the time of the stimulus and the time of the behavioral response but were free to blink after responding. Instructing participants to suppress blinks altogether is known to require cognitive effort (Lerner et al., 2009) and can impact ERPs (Ochoa & Polich, 2000). However, merely delaying the blinks until after the response seems to require less effort (although we know of no formal demonstrations of this). Instructions to delay blinks can be advantageous because they minimize the number of trials with blinks that might interfere with seeing the stimuli and reduce the possibility of differences in blink activity between conditions prior to the response. However, this procedure may also cause the blinks to be concentrated in the late part of the epoch. Differences in blinking across conditions may then be present late in the epoch, creating EOG confounds that might be mistaken for late ERP effects. The overall benefit of instructions to delay blinking will depend on the nature of a given study.

### 2.2. General artifact correction and rejection methods

Independent component analysis (ICA) was used for artifact correction. We used the EEGLAB runica() routine, which implements the infomax algorithm. This is probably the most widely used ICA algorithm for artifact correction (because it is the default in EEGLAB), and it is also one of the best-performing algorithms (Delorme et al., 2012). We used the default parameters, because our goal was to evaluate the effectiveness of a widely used approach rather than to determine the optimal approach.

The original ICA-based correction performed on the ERP CORE data did not follow recent recommendations for optimization (Dimigen, 2020; Klug & Gramann, 2021; Luck, 2022; Winkler et al., 2015). We therefore conducted a new ICA decomposition, in which we created a parallel dataset for each participant that was optimized for the decomposition. In this parallel dataset, we first applied a noncausal Butterworth band-pass filter with half-amplitude cutoffs at 1 Hz and 30 Hz and a roll-off of 12 dB/octave. Although this relatively narrow bandpass significantly distorts the time course of the ERP waveform (Zhang et al., 2023a, 2023b), it does not change the scalp distributions, so it does not interfere with the ICA decomposition process. However, it minimizes idiosyncratic noise that would otherwise degrade the decomposition. The data were then resampled at 100 Hz to increase the speed of the decomposition. Finally, we deleted break periods and time periods with non-biologically plausible outlier voltages from the continuous EEG, because these periods also contain idiosyncratic noise that would otherwise degrade the decomposition. This was achieved using the ERPLAB pop_erplabDeleteTimeSegments() function to delete break periods (defined as periods of at least 2 seconds without an event code) and the ERPLAB pop_continuousartdet() function to delete periods in which the peak-to-peak amplitude within a specific window length exceeded a threshold. The window length and threshold were set individually via visual inspection in the original ERP CORE resource, with a window length between 500 and 2000 ms and a threshold between 350 and 750 µV.

We included all EEG and EOG electrodes in the ICA decomposition except for the bipolar channels and any “bad” channels that required interpolation (as identified in the original ERP CORE dataset). Independent components (ICs) that represented blink artifacts were identified using ICLabel, an automatic IC classification system that was trained on a large number of datasets with manually labeled ICs (Pion-Tonachini et al., 2019). ICLabel assigns a probability that a given IC reflects a specific artifact. We classified an IC as reflecting blinks if ICLabel gave it a probability of at least 0.9 as reflecting blink activity. ICs with a lower probability are by definition more ambiguous, so we visually assessed the time-course match between these ICs and the VEOG-bipolar signal. A good match was occasionally observed for these ICs, and we also classified these ICs as reflecting blinks^4^. This semi-automatic approach to IC classification is quite common, but it involves the subjective judgment of experts and is therefore difficult to reproduce exactly. We therefore repeated all analyses using fully automatic classification (i.e., without visual inspection), once with a probability threshold of 0.9 and once with a probability threshold of 0.8. All three approaches yielded identical patterns of statistical significance for the analyses reported in this paper, except for one that is noted in the Results section. The specific results shown below came from the semi-automatic approach. Table 1 summarizes the number of ICs that were classified as reflecting blinks with each of these approaches.

**Table 1.** Mean number of blink-related independent components identified by three methods for each ERP component (range in parentheses)

|  | Fully automatic<br>classification<br>(probability = 0.8-1.0) | Fully automatic<br>classification<br>(probability = 0.9-1.0) | Semi-automatic<br>classification<br>(probability = 0.9-1.0 plus<br>visual inspection) |
| --- | --- | --- | --- |
| N400 | 1.40<br>(0-3) | 1.35<br>(0-3) | 1.48<br>(1-3) |
| ERN | 1.35<br>(1-2) | 1.13<br>(0-2) | 1.33<br>(1-2) |
| P3b | 1.10<br>(0-2) | 1.05<br>(0-2) | 1.13<br>(1-2) |
| MMN | 1.35<br>(0-2) | 1.18<br>(0-2) | 1.48<br>(1-2) |
| N170 | 1.38<br>(0-3) | 1.20<br>(0-2) | 1.33<br>(0-3) |

After the decomposition was performed, we transferred the component weights back to the original dataset that was not heavily filtered. The EEG was then reconstructed from the non-blink ICs. This procedure made it possible to obtain ICA weights using heavily filtered data that were not degraded by idiosyncratic noise, but the ICA weights were then applied to the original data so that the time course of the ERPs would not be distorted by the filtering.

Artifact detection and rejection were performed after artifact correction and after the data were epoched and baseline-corrected using the time periods shown in Table 2. Every epoch from every corrected EEG channel was subjected to two algorithms that detected somewhat different kinds of extreme values. We first applied ERPLAB’s simple voltage threshold algorithm with a threshold of 200 µV, which flagged any epochs in which the baseline-corrected voltage exceeded the threshold at any time point during the epoch. We then applied ERPLAB’s moving window peak-to-peak algorithm with a threshold of 100 µV, which computed the peak-to-peak amplitude within overlapping 200-ms windows across a given epoch. An epoch was flagged if the peak-to-peak amplitude within any window exceeded the threshold. These specific thresholds were chosen on the basis of an exploratory analysis comparing different threshold combinations, in which we found that these thresholds tended to yield the greatest overall reduction in noise. Details of this exploratory analysis can be found in the supplementary materials (Figures S2-S6). The epochs that were flagged for artifacts were then excluded when the averaged ERPs were computed. All channels were excluded for a given epoch if an artifact was flagged in any channel.

**Table 2.** Epoch window, baseline period, electrode site, and measurement window used for each ERP component.

|  | N400 | ERN | P3b | MMN | N170 |
| --- | --- | --- | --- | --- | --- |
| Epoch (ms) | -200 to 800 | -600 to 400 | -200 to 800 | -200 to 800 | -200 to 800 |
| Baseline Period (ms) | -200 to 0 | -400 to -200 | -200 to 0 | -200 to 0 | -200 to 0 |
| Measurement Channel | CPz | FCz | Pz | FCz | PO8 |
| Measurement Window (ms) | 300 to 500 | 0 to 100 | 300 to 600 | 125 to 225 | 110 to 150 |

Because the ICA-corrected data were used for the artifact detection process, blinks did not create extreme values and did not cause an epoch to be flagged for rejection. However, we wanted to count the number of trials with blinks in each condition to determine if blink rates differed across conditions. This was accomplished by applying ERPLAB’s step function routine to the uncorrected VEOG-bipolar channel. This routine slid a moving window across the epoch, and the absolute value of the difference between the mean voltage in the first half and second half of each window was computed (see Luck, 2014 for a detailed description). The largest of these values for a given epoch was then compared with a threshold. We used a window width of 200 ms, a step size of 10 ms, and a threshold of 100 µV.

In addition, one of our analyses excluded trials in which a blink occurred near the time of the stimulus, which prevents the perception of that stimulus. This was accomplished using the step function but limiting the time period to −200 to +200 ms.

### 2.3. Specific artifact correction and rejection approaches

We compared five different approaches to combining artifact correction with artifact rejection. In our baseline approach, labeled “*None*”, no artifact correction or rejection were applied. We simply epoched and baseline-corrected the EEG using the time windows specified in Table 2, and then averaged all the EEG epochs for a given condition except those with incorrect behavioral responses.

For the second approach, labeled “*ICA*”, we used the ICA approach described in the previous section to correct for blinks and then averaged all the EEG epochs except those with incorrect behavioral responses. No epochs were rejected because of EEG or EOG artifacts.

The third approach, labeled “*ICA+EV1*” was identical to the ICA approach except that epochs with extreme values (EVs) in any channel (as defined in the preceding section) were excluded during averaging. Note that the extreme values were assessed after artifact correction, so blinks did not lead to rejection.

The fourth approach, labeled “*ICA+EV1+Blink*”, was identical to the ICA+EV1 approach except that the averages also excluded epochs with blink activity around the time of the stimulus (as defined in the previous section). This excludes trials in which a blink would have prevented the perception of a visual stimulus.

The fifth approach, labeled “*ICA+EV2*”, was identical to the ICA+EV1 approach except that epochs were rejected only if extreme values were present in the measurement channel for a given component (listed in Table 2). By contrast, the ICA+EV1 approach excludes trials with extreme values in any channel. By narrowing extreme value detection to the measurement channel, we aimed to retain more trials.

We also explored two additional approaches in which channels exhibiting extreme values during an epoch were interpolated for that epoch rather than being rejected. This is not a common approach, and it did not work as well as rejection, so the results are described in the supplementary materials (see Figure S1).

### 2.4. Assessment of data quality

We computed the data quality separately for each of the five artifact minimization approaches across four scoring methods: mean amplitude, peak amplitude, 50% area latency, and peak latency. To estimate the noise, we focused on the difference waves of the specific components under investigation. In the case of the P3b component, for example, we computed a rare-minus-frequent difference wave, scored the amplitudes and latencies from this difference wave, and obtained the data quality for these scores. All analyses for a given component were restricted to the maximal channel for that component, as listed in Table 2.

Mean amplitude was defined as the mean voltage within the measurement window listed in Table 2. Peak amplitude was determined as the maximum positive voltage (for the P3b component) or the maximum negative voltage (for other components) within the measurement window, whereas peak latency was defined as the latency at which the peak amplitude occurred. To measure the 50% area latency, we computed the area bounded by the zero voltage line and the ERP difference wave during the measurement period (i.e., the area on the positive side of the zero line for the P3b and the area on the negative side of the zero line for the N170, MMN, N400, and ERN) and located the time point that divided the integral into equal halves. To enhance temporal precision, we upsampled the waveforms by a factor of 10 using spline interpolation before scoring the latencies (see Luck, 2014 for the rationale for upsampling and a more detailed description of the 50% area latency measure).

SME values were obtained individually from each participant for each score. For mean amplitude scores, we first computed the analytic SME (aSME) value for each parent waveform (e.g., the rare and frequent waveforms for the P3b component). Specifically, the mean amplitude score (i.e., the mean voltage across the measurement window) was obtained for each epoch for a given participant in a given condition (using the same epochs that are used for averaging), and the aSME was computed as the standard deviation of these scores divided by the square root of the number of epochs. We then computed the SME of the difference between waveforms as 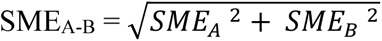, where SME_A-B_ is the SME of the difference between conditions A and B, and SME_A_ and SME_B_ are the SMEs of the two individual conditions (see Zhang et al., 2023a for details).

This approach is not valid for other scoring methods (e.g., peak amplitude, peak latency, 50% area latency). For these scores, we instead employed bootstrapping to estimate the SME (bootstrapped SME or bSME) directly from the relevant difference wave. The bootstrapping process is explained in detail in Luck et al. (2021) and requires simple scripting (example bSME scripts available at https://doi.org/10.18115/D58G91). In our study, we used 1,000 bootstrap iterations for each bSME value.

To obtain an aggregate measure of data quality across participants, we computed the root mean square (RMS) of the single-participant SME values (the *RMS(SME)*). This was computed by summing the squared single-participant SME values, dividing this sum by the number of participants, and then taking the square root. Note that we used the RMS across participants rather than the mean across participants because the RMS is more directly related to effect sizes and statistical power (Luck et al., 2021). Bootstrapping (with 10,000 iterations) was used to obtain the standard error of the RMS(SME) values.

## 3. Results

Each of the following sections presents the results for one of the five ERP components. The first section uses the N400 data to exemplify in detail our general method for assessing the effectiveness of artifact correction and rejection (see flowchart in Figure 1). This is followed by briefer sections on the ERN, P3b, MMN, and N170 components.

**Figure 1.**
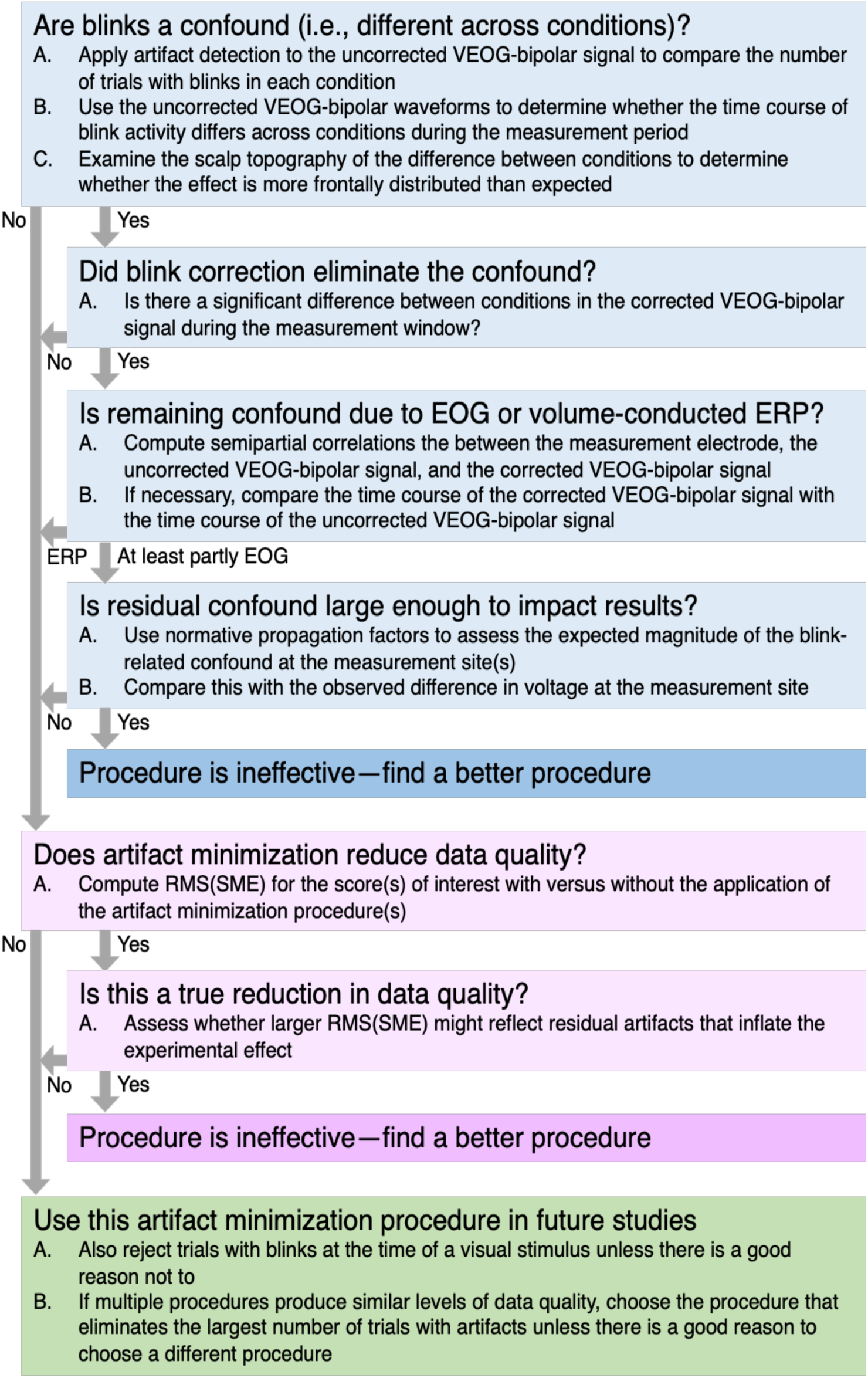
Summary of the present method for assessing the effectiveness of an artifact minimization approach for a given dataset.

### 3.1. Applying our general method to the N400 component

As shown in Figure 2a, the N400 component was elicited by means of a word-pair judgment paradigm. Each trial consisted of a red prime word followed by a green target word. Participants were tasked with indicating whether the target word was semantically related (p = 0.5, 60 trials) or unrelated (p = 0.5, 60 trials) to the preceding prime word by pressing one of two buttons. The N400 was measured from the unrelated-minus-related difference wave at the CPz electrode site.

**Figure 2.**
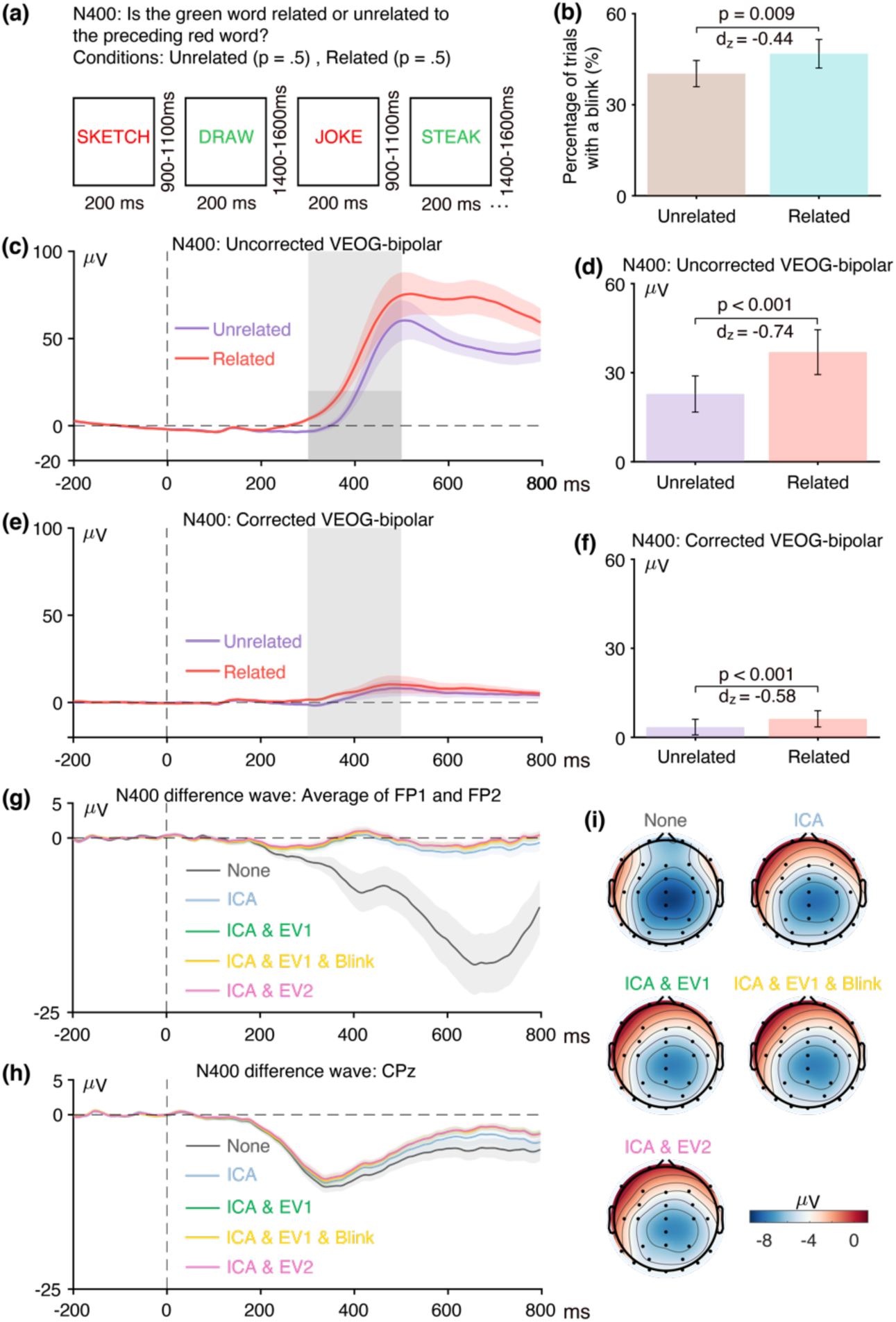
(a) N400 word pair judgment paradigm. (b) Percentage of trials with a blink for the parent waves, measured from uncorrected VEOG-bipolar channel. (c) Grand average ERP waveforms for semantically unrelated and semantically related targets in the uncorrected VEOG-bipolar electrode site. (d) Mean amplitudes from the uncorrected VEOG-bipolar channel during the N400 measurement window for the unrelated and related targets. (e) Grand average ERP waveforms for unrelated and related targets in the corrected VEOG-bipolar channel. (f) Mean amplitudes from the corrected VEOG-bipolar channel during the N400 measurement window for the unrelated and related targets. (g) Grand average ERP difference waves (unrelated minus related) for the five artifact minimization approaches at the FP1 and FP2 electrode sites. (h) Grand average difference waves for the five artifact minimization approaches at CPz. (i) Scalp maps of the mean amplitude measured from 300–500 ms in the grand average difference wave. Error bars show the standard error of the mean. The VEOG-bipolar signals were computed as upper minus lower.

#### 3.1.1 Assessment of eyeblink confounds in the N400 data

As shown in Figure 1, the first general goal of our method is to determine whether eyeblinks differed across conditions and were therefore a potential confound. The first step toward this goal is to examine the percentage of trials on which a blink occurred for the two conditions, as determined from the uncorrected VEOG-bipolar signal. As shown in Figure 2, blinks were significantly more common for the semantically related targets than for the semantically unrelated targets (t(39) = −2.77, p = 0.009, Cohen’s *d_z_* = −01.44). Thus, blink-related activity is a confound that could distort measurements of N400 activity.

As illustrated in Figure 2c, the next step is to examine the time course of the blink activity in the averaged ERPs from the uncorrected VEOG-bipolar channel (defined as the voltage above the eyes minus the voltage below the eyes). A very large deflection (>50 µV) was present, and it was larger for the semantically related targets than for the semantically unrelated targets from approximately 250 ms after stimulus onset through the end of the epoch. We measured the mean amplitude of this blink-related activity during the time window of the N400 component (300–500 ms). Consistent with the difference in blink frequency across conditions, we found that the mean amplitude was significantly greater for the semantically related targets than for the semantically unrelated targets (see Figure 2d; *t*(39) = −4.66, *p* < 0.001, Cohen’s *d_z_* = - 0.74). Note that this effect size of 0.74 was even greater than the effect size of 0.58 for the proportion of trials with blinks. In general, comparing the voltages across conditions (which is based on a continuous variable) is more sensitive than comparing the proportion of blinks (which is based on a categorical variable). In addition, the time course of the blink activity might differ across conditions even if the proportion of blinks does not differ.

The next step of our procedure is to examine how the blink confound, if present, propagates to the EEG electrodes. This is assessed by isolating the experimental effect with a difference wave, plotting this difference wave at key scalp electrodes, and making a scalp map of the difference wave during the measurement window. This is illustrated in panels g, h, and i of Figure 2, which show the unrelated-minus-related N400 difference wave for both the original data and the data following application of our artifact minimization approaches. At the FP1 and FP2 channels, where blink-related activity should be maximal, a very large negative deflection can be seen beginning at approximately 200 ms when no artifact correction was performed. This negativity presumably reflects the greater blink-related EOG activity observed for related targets than for unrelated targets in the uncorrected VEOG-bipolar signal. The voltage deflection in this N400 difference wave was negative-going because the difference wave was computed by subtracting the related trials (which had a larger positive VEOG deflection) from the unrelated trials (which had a smaller positive VEOG deflection).

Panel h shows the corresponding data from the CPz channel, which is the a priori measurement site for the N400. The difference wave was again more negative when no artifact correction was performed than when our artifact minimization procedures were performed. However, the difference between the waveforms with and without artifact minimization was much smaller at CPz than at FP1 and FP2, reflecting the fact that only a small percentage of the blink-related activity propagates to CPz. The blink-related activity that was present without artifact minimization can also be observed in the scalp maps of the unrelated-minus-related difference (panel i). Specifically, the topography of the difference was more frontal when blink correction was not performed.

Together, these results demonstrate that blink-related activity was greater for semantically related targets than for semantically unrelated targets and was therefore a potential confound in comparing these two conditions. If this activity is not reduced to negligible levels, any differences in apparent ERP activity between the semantically related and unrelated targets after approximately 200 ms could reflect differences in blinking rather than differences in the N400 component. Thus, some method is necessary to minimize the blink-related artifacts.

#### 3.1.2. Effectiveness of ICA at minimizing eyeblink confounds in the N400 data

Given that a significant confound was present in the uncorrected VEOG-bipolar signal, the next step is to determine whether this confound was eliminated in the ICA-corrected data. This involves using the ICA-corrected data to create a *corrected VEOG-bipolar* channel (computed as the corrected F2 channel minus the corrected VEOG-lower channel). As shown in Figure 2e, most of the blink activity was removed, and most of the difference in blink-related activity between semantically related and semantically unrelated targets was also eliminated. However, the corrected VEOG-bipolar signal was still 2.79 µV greater for related targets than for unrelated targets during the N400 measurement window (t(39) = −3.70, p < 0.001, Cohen’s *d_z_* = −0.58). This raises the possibility that the ICA-based correction did not fully eliminate the confounding EOG activity.

However, it is also possible that the difference between conditions in the uncorrected VEOG-bipolar waveform does not reflect blink activity but instead reflects volume-conducted N400 activity. That is, if the N400 activity is larger (more negative) above the eyes than below the eyes, then this will add a negative voltage to the above-minus-below subtraction used to create the VEOG-bipolar channel. And because the N400 was larger for unrelated targets than for related targets, this added voltage in the VEOG-bipolar channel should be larger (more negative) for unrelated targets than for related targets. This is exactly what was observed, with a more negative VEOG-bipolar voltage for the unrelated targets than for the related targets (see Figure 2e). Thus, it is not obvious whether the remaining difference between related and unrelated targets in the corrected VEOG-bipolar channel reflects volume-conducted N400 activity or eyeblink activity that was not fully eliminated by ICA.

To assess this possibility, we used semipartial correlations to test for the presence of volume-conducted ERP activity in the corrected VEOG-bipolar signal^5^. Specifically, we computed the correlation between the corrected unrelated-minus-related difference voltage at the CPz electrode and the corrected unrelated-minus-related difference voltage in the VEOG-bipolar channel after partialling out variance explained by the unrelated-minus-related difference voltage in the uncorrected VEOG-bipolar channel. This approach is based on the assumption that the voltage at CPz and the voltage in the uncorrected VEOG-bipolar signal will share some variance due to the propagation of EOG and ERP activity between them, and the simple correlation between CPz and the corrected VEOG-bipolar signal could reflect residual EOG activity that propagated to CPz. However, if we partial out variance due to the uncorrected VEOG-bipolar signal (which is dominated by true EOG activity), then any remaining correlation between CPz and the corrected VEOG-bipolar signal should almost entirely reflect the propagation of the N400 to the VEOG-bipolar channel^6^. Because we are mainly interested in activity that might confound the N400 effect, we used the mean voltage during the N400 measurement window (see Table 2), measured from the unrelated-minus-related difference wave. We did not find a significant semipartial correlation between the corrected CPz signal and the corrected VEOG-bipolar signal (r(38) = −0.026, p = .876), providing no evidence that the difference between related and unrelated trials in the corrected VEOG-bipolar signal could be explained by volume-conducted N400 activity.

An analogous approach can be used to assess whether the corrected VEOG-bipolar signal contains residual EOG activity. In this approach, the correlation between the uncorrected and corrected VEOG-bipolar signals is computed after partialing out variance explained by the measurement electrode (again using the mean voltage of the difference wave during the measurement period). In the N400 data, we found a significant semipartial correlation between the uncorrected and corrected VEOG-bipolar signals (r(38) = .397, p = .012). This correlation provides evidence that at least some of the variance in the unrelated-minus-related difference wave in the corrected VEOG-bipolar signal could be explained by eyeblink activity, indicating that ICA did not completely eliminate the confounding effects of eyeblinks. Consistent with this conclusion, the corrected VEOG-bipolar signal contained a noticeable deflection for both the unrelated and related trials that approximately matched the blink-related deflection in the uncorrected VEOG-bipolar signal.

Note that the *lack* of a significant semipartial correlation between the uncorrected and corrected VEOG-bipolar signals cannot be used as evidence that ICA was effective at minimizing blink-related confounds. This is because the blink-related activity may be responsible for a large portion of the shared variance that is being partialed out^7^. We will discuss this in more detail in the case of the ERN.

When there is reason to believe that ICA was not completely effective at removing the confound in the corrected VEOG-bipolar signal, the next step is to determine whether the residual confound was large enough to have a meaningful impact when propagated to the measurement electrode. That is, once the 2.79 µV unrelated-minus-related difference that was measured in the VEOG-corrected channel propagates to the scalp electrodes, how much would it impact the uncorrected-minus-corrected N400 effect measured from the CPz channel? The normative values provided by Lins et al. (Lins et al., 1993) indicate that we would expect a propagated voltage of approximately 19% at Fz, approximately 10% at Cz, and approximately 5% at Pz. Propagation values are unavailable for CPz but are presumably between those at Cz and Pz. The 2.79 µV effect in the corrected bipolar VEOG channel would therefore be expected to produce an effect of approximately 0.53 µV at Fz, 0.28 µV at Cz, and 0.14 µV at Pz, with a value somewhere between 0.14 and 0.28 µV at CPz. This would be less than 4% of the measured difference between related and unrelated trials of −8.11 µV in the N400 measured at CPz. Thus, although ICA did not completely eliminate the confounding eyeblink activity at CPz, the residual activity was negligible for the purposes of this specific study.

However, the confounding effects on N400 measurements might be larger in other paradigms and participant populations, and even a small confound in the measurement electrode might be problematic for answering some scientific questions (e.g., when the confound is large relative to the size of the effect at the measurement electrode). The present results therefore indicate that researchers should be very cautious about ocular confounds in N400 studies if they are using ICA-based artifact correction. In addition, researchers should assess whether confounds are present in their own data and consider applying our method (as summarized in Figure 1) for assessing whether the confound was reduced to negligible levels.

#### 3.1.3. Alternative ICA decomposition of the N400 data

We also considered the possibility that the residual activity in the corrected VEOG-bipolar channel was a result of the specific ICA decomposition approach used in the present analyses, which involved aggressive filtering of low frequencies and the use of the automated ICLabel routine to determine which ICs represented artifacts (along with manual confirmation). To assess this possibility, we performed the same analysis using the corrected data from the ERP CORE resource, in which the ICA decomposition did not involve aggressive high-pass filtering and the artifact-related ICs were selected “by eye” on the basis of their scalp maps and time courses.

We again found a statistically significant difference of 4.01 µV between related and unrelated targets during the N400 latency window in the corrected VEOG-bipolar channel (t(39) = 6.00, p <0.001, Cohen’s *d_z_* = 0.95). We also found a significant semipartial correlation between the uncorrected VEOG-bipolar channel and the corrected VEOG-bipolar channel (r(38) = 0.397, p = 0.012) whereas the semipartial correlation between the CPz channel and the corrected VEOG-bipolar channel was again not significant (r(38) = −0.024. p = 0.876). Thus, the apparent failure of ICA to completely eliminate blink-related confounds generalized to a different ICA decomposition. Of course, ICA might be more effective with different parameters, and there may be other artifact correction approaches that are more effective (e.g., second-order blind inference or source-space reconstruction; Jonmohamadi et al., 2014; Joyce et al., 2004). However, the goal of the present study was to determine whether the commonly-used approach implemented here is effective, not to answer the much more challenging question of which correction approach is best.

#### 3.1.4 Assessment of data quality in the N400 data

Once it has been demonstrated that artifact correction has reduced any blink-related confounds to negligible levels, the next step is to assess whether the combination of correction and rejection increases or decreases the data quality (see Figure 1). In this section, we therefore evaluate the effectiveness of our artifact correction and rejection approaches in reducing the RMS(SME).

Figure 3 shows the RMS(SME) values at the a priori measurement site (CPz) for each combination of the four scoring methods and the five artifact minimization approaches. Interestingly, the ICA-only method did not lead to lower RMS(SME) values (i.e., did not lead to improved data quality). This is presumably because such a small proportion of blink activity is propagated to the CPz measurement site that blinks do not meaningfully increase trial-to-trial variability at this site. This result is consistent with Delorme (2023), who found that ICA-based artifact correction did not increase the ability to detect significant effects in several datasets.

**Figure 3.**
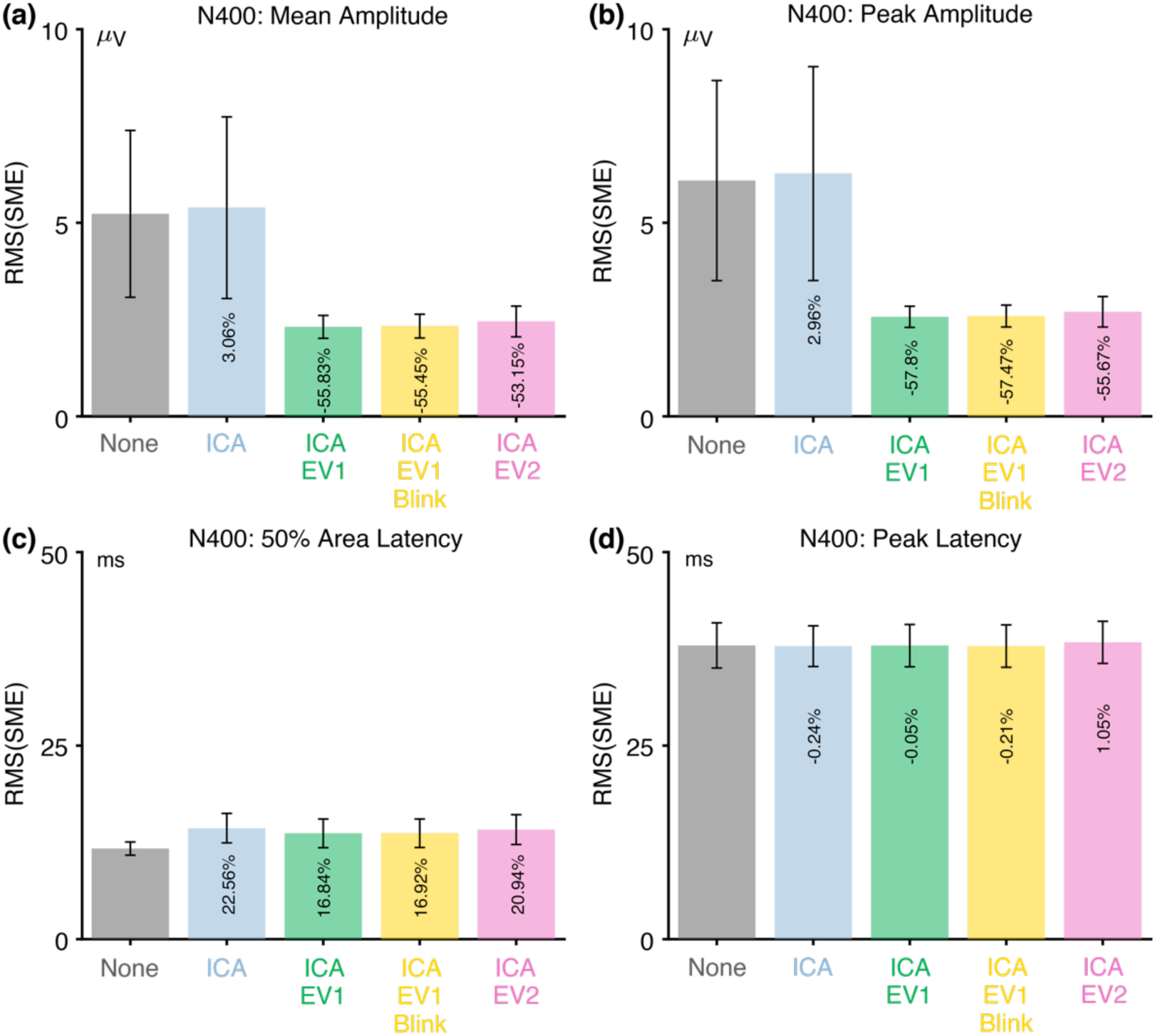
Root mean square of the standardized measurement error (RMS(SME)) from the N400 experiment for four different scoring methods and five different artifact minimization approaches. Smaller RMS(SME) values indicate higher data quality (less noise). Error bars show the standard error of the RMS(SME) values.

However, the three approaches that included rejection of trials with extreme values led to much better data quality for the two amplitude scoring methods compared to no artifact minimization and compared to the ICA-only approach. Indeed, rejecting trials with extreme values reduced the RMS(SME) by more than 50%. In other words, although artifact rejection reduced the number of trials, the net effect was still an improvement in data quality. For peak latency, the RMS(SME) was similar across all five approaches.

For 50% area latency, the RMS(SME) was actually slightly worse (larger) for all of the approaches that included ICA-based correction compared to the data with no artifact minimization. However, this may be a consequence of the fact that the blink-related activity increased the size of the unrelated-minus-related difference wave (as shown in Figure 2h). In other words, if the difference wave is larger as the result of greater blink activity for related targets than for unrelated targets, then the 50% area latency can be measured more consistently. However, the resulting latency values will be distorted by the blink-related activity, so the values might be misleading even if they are more consistent. Thus, although the no-minimization approach might appear to be advantageous for scoring the 50% area latency, this advantage is illusory. This illustrates the importance of assessing confounds in addition to data quality.

Table 3 shows the proportion of trials that were excluded because of artifacts for the three approaches that involved artifact rejection (ICA+EV1, ICA+EV1+Blinks, ICA+EV2). There were very few trials with blinks at the time of the stimulus in the N400 experiment, so the percentage of rejected trials was only slightly greater for the ICA+EV1+Blinks approach than for the ICA+EV1 approach. More trials were rejected for these approaches than for the ICA+EV2 approach, in which trials were rejected only if the extreme values occurred in the measurement channel (CPz). However, fewer than 5% of trials were rejected for any of these approaches. In addition, these three approaches led to similar RMS(SME) values.

**Table 3.**
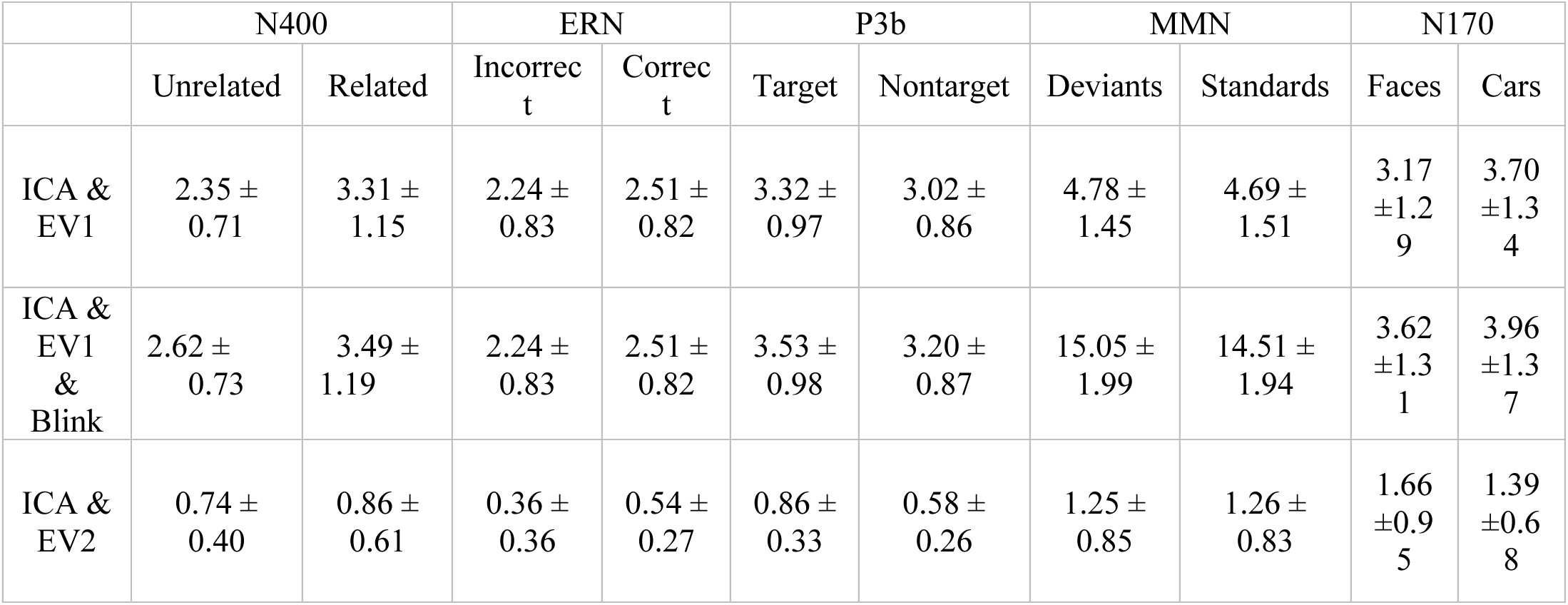
Percentage of rejected trials for each condition of each ERP component (in %).

Participants in the original ERP CORE study were excluded from the final analyses if more than 25% of trials were rejected because of extreme values. The effects of excluding participants are complicated because exclusion impacts the degrees of freedom as well as the data quality, so we are not excluding participants with excessive rejection in the main analyses of the present study. However, because excluding participants with large numbers of artifacts is a common procedure, we have provided supplementary analyses showing the RMS(SME) values when these participants were excluded (see supplementary Table S1 and Figures S7-S11). One participant exceeded our threshold for exclusion for the EV1 approach in the N400 paradigm. As shown in Figure S7, excluding this participant further improved the RMS(SME) values for the amplitude scores, with little impact on the RMS(SME) values for the latency scores.

#### 3.1.5 Recommendations for the N400

For studies like the ERP CORE N400 experiment, the present results indicate that blinks are a potential confound that can contribute to differences between conditions, even in the central and parietal channels where the N400 is typically scored. Thus, some method for minimizing the impact of blink-related activity (e.g., artifact correction or rejection) is essential in N400 experiments.

ICA-based blink correction substantially reduced the blink artifact in the present analyses, but a statistically significant artifactual difference between conditions remained in the corrected VEOG-bipolar channel. After propagating to the central and parietal channels, the residual artifactual difference between conditions was negligible (less than half a microvolt). However, an effect of this size might still be large enough to produce an incorrect interpretation of the results in some studies, and larger blink artifacts may be present in other paradigms or participant populations. We therefore recommend that all N400 studies quantify the difference between conditions in the corrected VEOG-bipolar signal. If the difference is significant, the propagation factors provided by Lins et al. (1993) can be used to determine whether the residual EOG activity is large enough to meaningfully impact the results in the channels where the N400 is being measured.

We also found that excluding trials with extreme values improved the data quality for the amplitude scores, with the reduction in number of trials being outweighed by the reduction in noise. There were no meaningful differences in data quality between the three rejection approaches examined here. We therefore recommend the EV1 approach to extreme values, which ensures that the data are clean in all channels (which may be important for scalp maps and other analyses). However, there may be studies in which it is advantageous to reject trials only when the extreme values occur in the measurement channel. In addition, we recommend rejecting trials with blinks at the time of the stimulus that might interfere with the perception of the stimulus unless there is a good reason not to. Thus, we recommend the ICA+EV1+Blinks approach for studies like the ERP CORE N400 experiment.

These recommendations, along with a summary of the key N400 results, are provided in Table 4.

**Table 4.** Summary of results and recommendations.

|  | Significant confound in VEOG-bipolar signal |  | Significant semipartial correlation with corrected VEOG |  | Recommendation for artifact minimization |
| --- | --- | --- | --- | --- | --- |
|  | Before correction | After correction | Uncorrected VEOG | Measured Channel |  |
| N400 | ✓ | ✓ | ✓ | × | ICA + EV1 + Blink |
| ERN | ✓ | ✓ | × | ✓ | ICA + EV1* + Blink |
| P3b | ✓ | × | ✓ | × | ICA + EV1 + Blink |
| MMN | × | × | ✓ | × | ICA* + EV1 |
| N170 | ✓ | ✓ | × | ✓ | ICA + EV1 + Blink |

ICA: Correction of blinks using independent component analysis; EV1: Rejection of trials with extreme values in any channel after artifact correction; Blink: Rejection of trials with blinks within $\pm 200$ ms of stimulus onset.
\* Optional

#### 3.1.6 When to assess the effectiveness of artifact minimization

Ideally, the effectiveness of artifact minimization would be assessed in an a priori manner. That is, it would be applied to one or more previous datasets, and the results would then be used to determine which minimization approach to use in future datasets. It is potentially problematic to use a given dataset to determine the best artifact minimization approach and then use that approach to analyze the same dataset. However, it is not clear how this post hoc approach could actually bias the results and increase the Type I error rate, because our method for assessing artifact minimization approaches does not depend on the extent to which differences in brain activity are present. On the other hand, using the same dataset twice in this manner might lead to non-obvious biases. Thus, it would be safest to use previous datasets to determine whether a given artifact minimization approach is adequate and then use this approach in an a priori manner for future studies.

### 3.2 The error-related negativity (ERN)

As illustrated in Figure 4a, we used a flankers paradigm to elicit the ERN. Each display consisted of a central arrow accompanied by four flanking arrows, which pointed in the same direction (p = .5) or opposite direction (p = .5) as the central arrow. Participants were tasked with determining whether the central arrow pointed leftward (p = .5, 200 trials) or rightward (p = .5, 200 trials) and responded by pressing a button with the corresponding hand. Each participant made correct responses on 352.1 ± 36.6 trials and incorrect responses on 42.0 ± 22.6 trials (except one participant, who had only 2 incorrect responses and was excluded from all analyses). The ERN was measured from the incorrect-minus-correct difference wave at the FCz electrode site.

**Figure 4.**
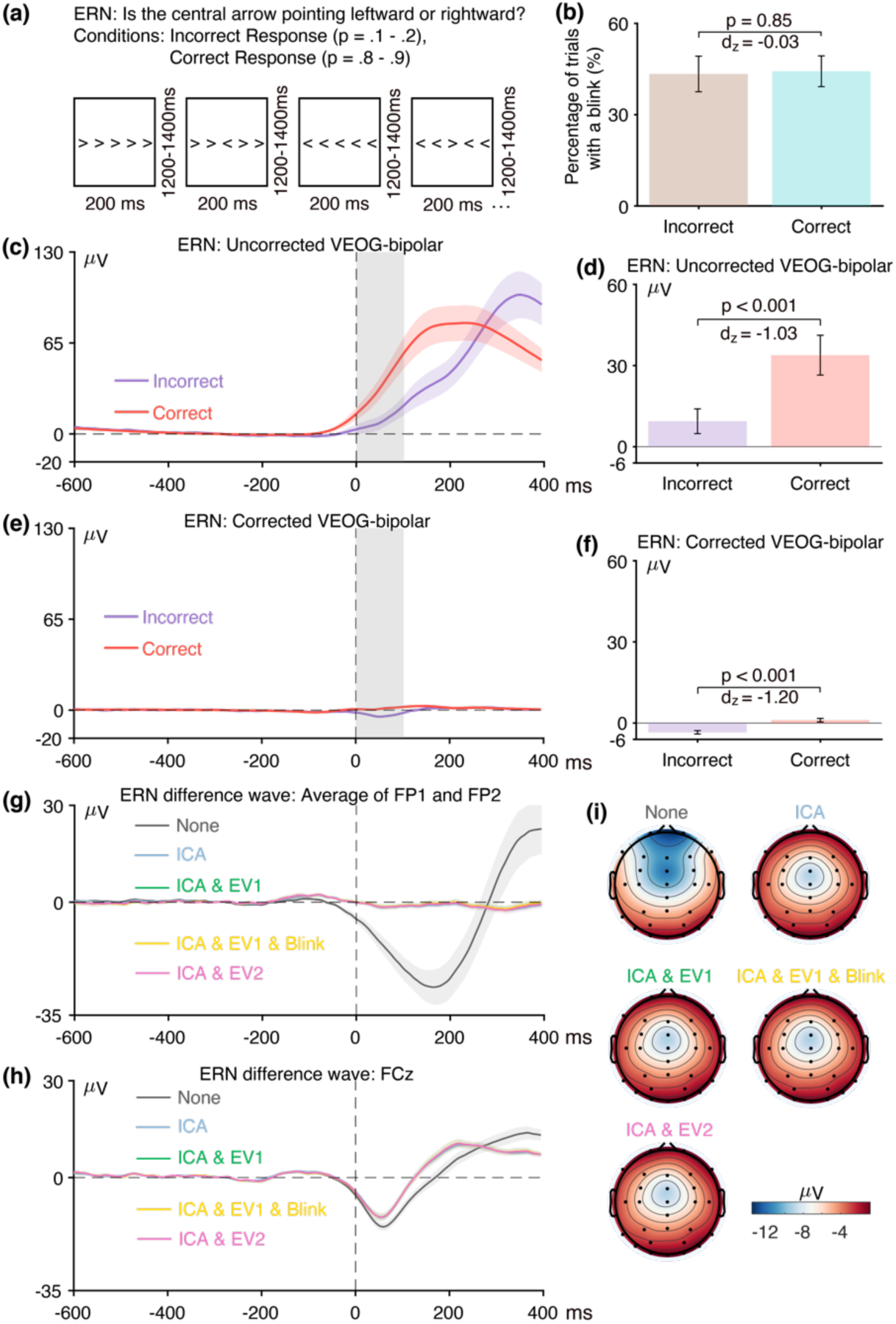
(a) Flankers task used to elicit the error-related negativity (ERN). (b) Percentage of trials with a blink for the parent waves, measured from uncorrected VEOG-bipolar channel. (c) Grand average ERP waveforms for the incorrect and correct conditions in the uncorrected VEOG-bipolar electrode site. (d) Mean amplitudes from the uncorrected VEOG-bipolar channel during the ERN measurement window for incorrect and correct trials. (e) Grand average ERP waveforms for the incorrect and correct conditions in the corrected VEOG-bipolar channel. (f) Mean amplitudes from the corrected VEOG-bipolar channel during the ERN measurement window for the incorrect and correct trials. (g) Grand average ERP difference waves (incorrect minus correct) for the five artifact minimization approaches at the FP1 and FP2 electrode sites. (h) Grand average difference waves for the five artifact minimization approaches at FCz. (i) Scalp maps of the mean amplitude measured from 0–100 ms in the grand average difference wave. Error bars show the standard error of the mean. The VEOG-bipolar signals were computed as upper minus lower. Note that the ERN data were response-locked rather than stimulus-locked, so time zero is the time of the response.

#### 3.2.1 Assessment of eyeblink confounds in the ERN data

The first step was to determine whether eyeblinks differed across conditions and were therefore a potential confound. Figure 4b shows the percentage of trials on which a blink occurred for the incorrect versus correct responses, as determined from the uncorrected VEOG-bipolar channel. Blinks were approximately equally likely on both correct trials and error trials (t(38) = −0.19, p = 0.85, Cohen’s *d_z_* = −0.03).

Figure 4c shows the grand average waveforms from the uncorrected VEOG-bipolar channel (upper minus lower). A very large deflection (>50 µV) was present, beginning just before the time of the response. This deflection was larger for correct responses than for incorrect responses from approximately −50 ms to 300 ms, and was then larger for incorrect responses than for correct responses until the end of epoch. It was significantly more positive for correct responses than for incorrect responses during the ERN measurement window (0–100 ms, Figure 4d; t(38) = −6.42, p < 0.001, Cohen’s *d_z_* = −1.03). Thus, even though the likelihood of a blink did not differ between incorrect and correct trials, the time course differed, creating a potential confound during the ERN measurement window. This demonstrates the importance of examining the time course of blink activity in averaged EOG waveforms and not just the likelihood of blinks (see Stern et al., 1984).

Panels g, h, and i of Figure 4 show how the blink confound—as assessed in the incorrect-minus-correct difference wave—propagated to the scalp ERPs. At the FP1 and FP2 channels, where blink-related activity should be maximal (panel g), the uncorrected waveform shows a large negative deflection followed by a large positive deflection. This is exactly what would be expected from the uncorrected VEOG-bipolar signal (panel c), in which the voltage was initially more positive for the correct trials and then became more positive for the incorrect trials. That is, the initial greater positive voltage for the correct trials relative to the incorrect trials created an initial negative voltage when the correct trials were subtracted from the incorrect trials in the difference wave; the subsequent greater positive voltage for the incorrect trials relative to correct trials in the VEOG created a late positive voltage in the incorrect-minus-correct difference wave. Note that the pattern produced by the blink activity in the difference wave at FP1 and FP2 is qualitatively similar to the pattern expected from true brain activity on the basis of prior research, consisting of an initial negative ERN followed by a later positive Pe. These deflections were greatly reduced in the waveforms after ICA-based blink correction was performed.

Panel h shows the corresponding data from the ERN measurement site, FCz. When artifact correction was performed, the typical pattern of an ERN followed by a Pe was observed, with a transition from negative to positive at approximately 130 ms. Both the negative and positive peaks were larger without artifact correction, and the negative-positive transition was shifted to approximately 200 ms. This is exactly what would be expected if true ERN and Pe deflections were present in the corrected data, with a volume-conducted EOG artifact summing with these effects in the uncorrected data. The residual artifact was quite large at FCz, both because the difference in blink activity between correct and incorrect trials was large and because FCz is fairly close to the source of the artifact. The blink-related activity that was present without artifact minimization can also be observed in the scalp maps of the incorrect-minus-correct difference (Figure 4i). Specifically, the topography of the difference was more frontal when the blink-related activity was not minimized.

Together, these results demonstrate that blinks are a very worrisome potential confound in ERN experiments. Specifically, the eyeblink confound would be expected to produce a more negative voltage for error trials than for correct trials at FCz immediately after the response (just like the ERN) followed by a more positive voltage for error trials than for correct trials (just like the Pe). Thus, if eyeblink confounds are not completely eliminated, they could easily masquerade as, or artificially augment, the ERN and Pe effects. Consequently, additional work is needed to be certain that the ERN and Pe effects observed after correction are not volume-conducted EOG artifacts.

#### 3.2.2. Effectiveness of ICA at minimizing eyeblink confounds in the ERN data

Figure 4e shows the grand average waveforms for the corrected VEOG-bipolar channel. Most of the blink activity was eliminated, and most of the difference in blink-related activity between the incorrect and correct trials was also eliminated. However, there was still a 4.51 µV difference between the incorrect and correct trials in the corrected VEOG-bipolar channel during the ERN measurement window, which was statistically significant (t(38) = −7.49, p < 0.001, Cohen’s d_z_ = −1.20). This raises the possibility that the ICA-based correction did not fully eliminate the confounding EOG activity, just as was observed in the N400 data.

However, this effect may instead reflect volume-conducted ERN activity. That is, if the difference in ERN voltage between incorrect and correct trials is larger (more negative) above the eyes than below the eyes, then this will appear as a more negative voltage for correct trials than for incorrect trials in the VEOG-bipolar channel. We used our semipartial correlation approach to determine whether the difference between the incorrect and correct trials in the corrected VEOG-bipolar channel reflects a failure of correction or volume-conducted ERN activity. That is, we quantified the extent to which variance in the corrected VEOG-bipolar channel can be uniquely explained by variance in the uncorrected VEOG-bipolar channel (which primarily contains blink activity) and by variance in the FCz channel (which primarily contains ERN activity). The correlations were computed using the voltage from 0-100 ms measured from the single-participant incorrect-minus-correct difference waves.

We found that the semipartial correlation between the FCz channel and the corrected VEOG-bipolar channel was substantial and statistically significant (r(37) = 0.43, p = 0.007)^8^. This indicates that at least some of the difference between incorrect and correct trials in the corrected VEOG-bipolar signal reflects volume-propagated ERN activity. By contrast, the semipartial correlation between the uncorrected and corrected VEOG-bipolar signals during the ERN measurement window was relatively small and not statistically significant (r(37) = 0.248, p = 0.134).

However, the lack of a significant semipartial correlation between the uncorrected and corrected VEOG-bipolar channels is not sufficient to conclude that there is no residual EOG activity in the corrected data. First, this would rely on accepting the null hypothesis. Second, some of the blink activity will produce shared variance between the uncorrected VEOG-bipolar signal and the signal at the measurement electrode. This shared blink activity is not captured by the semipartial correlation. To take an extreme example, imagine that FP2 was used as the measurement electrode. If the activity at FP2—which is largely blink activity—was partialed out from the correlation between the uncorrected and uncorrected VEOG-bipolar signals, this would remove almost all of the blink-related variance. The remaining correlation would therefore provide little information about whether similar blink activity was present in the uncorrected and corrected VEOG-bipolar signals. Thus, whereas the *presence* of a significant semipartial correlation between the uncorrected and corrected VEOG-bipolar signals provides good reason to believe that residual EOG activity is present after correction, the *absence* of a significant correlation is not sufficient to conclude that correction was successful.

To provide additional evidence about the presence of residual EOG activity after correction, we examined the time course of the corrected and uncorrected VEOG-bipolar signals. The ICA blink correction procedure uses the same unmixing and mixing matrices at all time points, and the scalp distribution of the true blink activity is also constant across time points. As a result, any residual EOG activity in the corrected EOG-bipolar signal should have approximately the same time course as the uncorrected VEOG-bipolar signal. If the difference between conditions in the corrected VEOG-bipolar signal is limited to the time at which the experimental effect is clearly present in the corrected data from the measurement electrode, whereas the difference between conditions in the uncorrected VEOG-bipolar signal extends more broadly, this provides good evidence that the correction procedure was successful and little or no residual EOG activity was present to confound the comparison of the conditions.

In the ERN data, for example, the difference between correct and incorrect trials in the corrected VEOG-bipolar signal was limited to the time period of the ERN component (ca. 0-150 ms; see Figure 4e). By contrast, there were large differences between correct and incorrect trials in the uncorrected VEOG-bipolar signal between 300 and 400 ms. If residual EOG activity were present in the corrected VEOG-bipolar signal, it would have been visible in this late period. We can therefore conclude that our implementation of ICA-based artifact correction was successful at minimizing the confounding blink activity for the ERN in this particular study.

#### 3.2.3 Assessment of data quality in the ERN data

Figure 5 shows the RMS(SME) values at the a priori measurement site (FCz) for each combination of the four scoring methods and the five artifact minimization approaches for ERN. For the two amplitude scoring methods, the RMS(SME) values were slightly better (smaller) for the four artifact minimization approaches (all of which included ICA-based blink correction) than when no artifact minimization was performed. Excluding trials with extreme values reduced the RMS(SME) slightly relative to the ICA-only approach. These results are again consistent with the finding of Delorme (2023) that ICA-based artifact correction did not increase the ability to detect significant effects.

**Figure 5.**
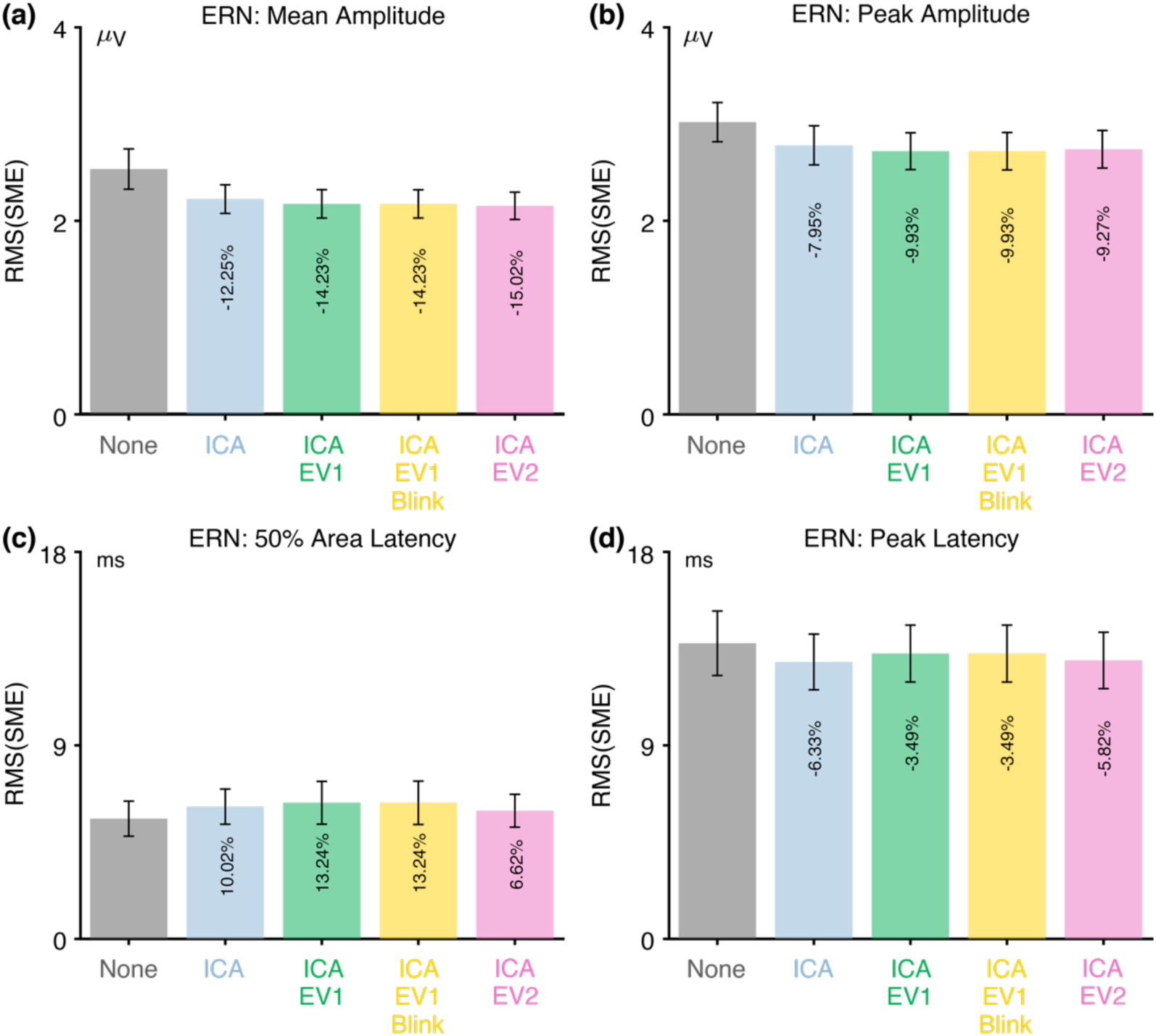
Root mean square of the standardized measurement error (RMS(SME)) from the ERN experiment for four different scoring methods and five different artifact minimization approaches. Smaller RMS(SME) values indicate higher data quality (less noise). Error bars show the standard error of the RMS(SME) values.

There was very little impact of any of the artifact minimization approaches for peak latency. For 50% area latency, however, the RMS(SME) was actually slightly better (smaller) for the no-minimization approach than for the other approaches. However, this may be a consequence of the fact that the blink-related activity increased the size of the incorrect-minus-correct difference wave (as shown in Figure 4h). In other words, if the difference wave is larger, then the 50% area latency can be measured more consistently. However, the resulting latency values will be distorted by the blink-related activity, so the values might be misleading even if they are measured more precisely.

Table 3 shows that fewer than 3% of trials were rejected for any of the three approaches that involved artifact rejection (ICA+EV1, ICA+EV1+Blinks, ICA+EV2).

#### 3.2.4 Recommendations for the ERN

For studies like the ERP CORE ERN experiment, the present results indicate that blinks are a particularly worrisome confound, because they may produce the same negative-positive sequence of voltages on the scalp as the ERN and Pe components. Thus, significant care is necessary to make sure that the artifact minimization procedure is effective. For example, if ERN activity is compared across groups, and the ERN appears to be larger in one group than in another, it would be essential to demonstrate that this is not a consequence of differences in blink-related activity.

Fortunately, we found that ICA-based blink correction did an excellent job of eliminating blink-related confounds. Although there was a substantial and statistically significant difference between correct trials and error trials in the corrected VEOG-bipolar signal, this difference appeared to primarily reflect volume-conducted ERN activity rather than uncorrected blink activity. However, ICA may not work this well in all datasets. For example, ICA may not remove blink-related confounds as well in studies with different numbers of electrodes, a shorter period of data, noisier data, and so on. We would therefore recommend using ICA-based blink correction in ERN experiments but carefully assessing its effectiveness rather than simply assuming that it completely eliminated blink-related confounds.

If evidence of a confound remains, the propagation factors provided by Lins et al. (1993) can be used to determine whether the residual EOG activity is large enough to meaningfully impact the results in the channels where the ERN is being measured.

We found that ICA-based artifact correction produced a modest improvement in data quality for amplitude scores. However, excluding trials with extreme values had minimal additional impact, indicating that the reduction in the number of trials produced by artifact rejection was approximately equally balanced by the reduction in noise. Given that the rejection of trials with extreme values did not hurt the data quality, and that it is presumably better not to include trials with extreme values if there is no cost for excluding them, we recommend rejecting trials with extreme values in datasets like the ERP CORE ERN experiment. However, very few trials were rejected in the present dataset, and rejection might have a larger effect in noisier datasets. In such datasets, it would be worthwhile to compute RMS(SME) values with and without rejection to determine whether the reduction in the number of trials resulting from artifact rejection is outweighed by the reduction in noise. In any case, trials in which the eyes were closed at the time of the stimulus should ordinarily be excluded. These recommendations, along with a summary of the key ERN results, are provided in Table 4.

### 3.3 The P3b component

Figure 6a illustrates the active visual oddball task that was used to elicit the P3b component. Participants were presented with a random sequence of five letters (A, B, C, D, and E), each with an equal probability (0.2). In every block, one specific letter was assigned as the target, and participants were instructed to press a designated button when the target letter appeared and a different button for any non-target letter. For example, if C was defined as the target, participants were instructed to press the target button when C appeared and the non-target button for the letters A, B, D, or E. Each letter served as the target in one block of trials. Over the experiment, each participant encountered 40 target trials and 160 non-target trials. The P3b was measured from the target-minus-nontarget difference wave at the Pz electrode site.

**Figure 6.**
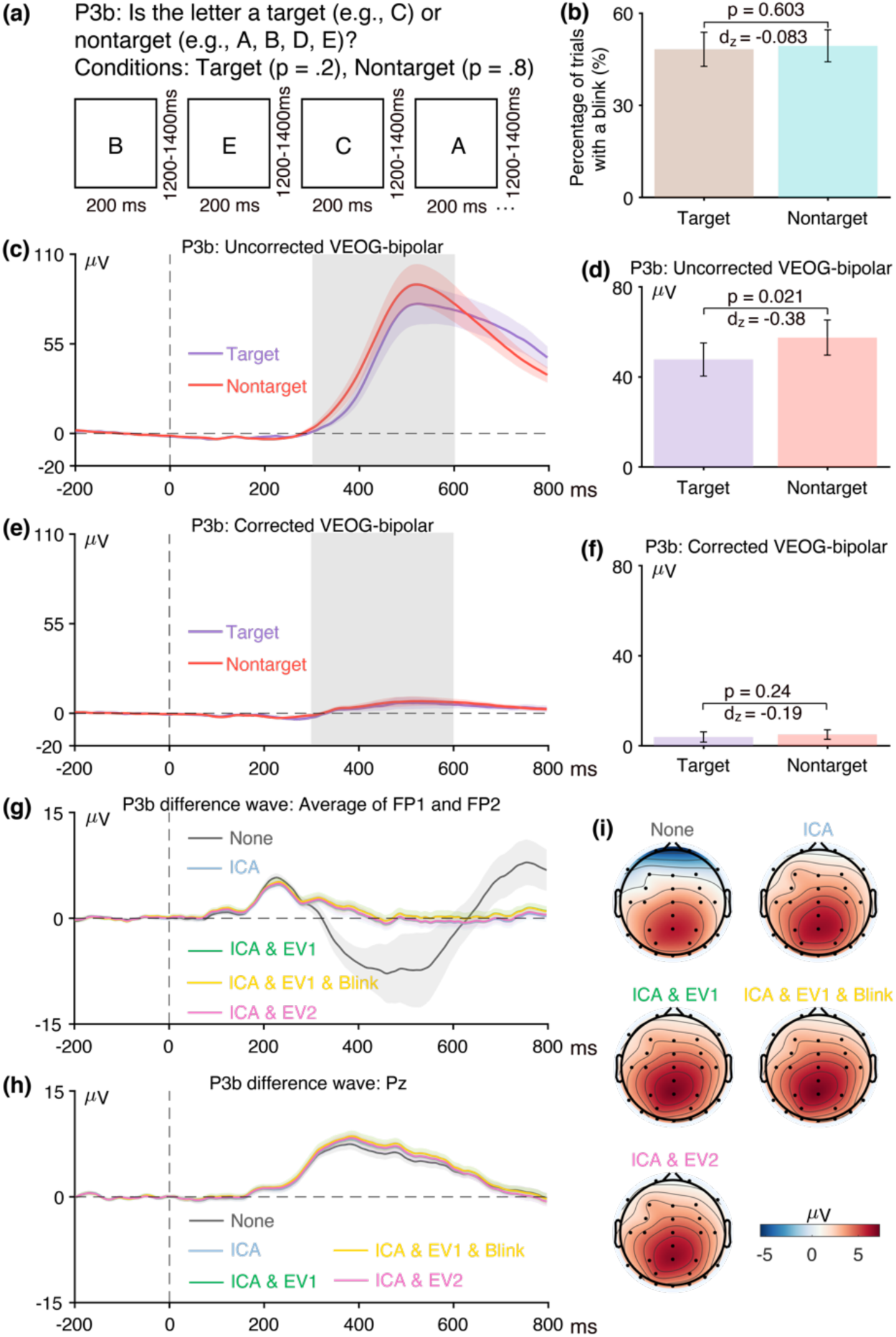
(a) Active visual oddball paradigm used to elicit the P3b component. (b) Percentage of trials with a blink for the parent waves, measured from uncorrected VEOG-bipolar channel. (c) Grand average ERP waveforms for targets and nontargets in the uncorrected VEOG-bipolar electrode site. (d) Mean amplitudes from the uncorrected VEOG-bipolar channel during the P3b measurement window for targets and nontargets. (e) Grand average ERP waveforms for targets and nontargets in the corrected VEOG-bipolar channel. (f) Mean amplitudes from the corrected VEOG-bipolar channel during the P3b measurement window for targets and nontargets. (g) Grand average ERP difference waves (target minus nontarget) for the five artifact minimization approaches at the FP1 and FP2 electrode sites. (h) Grand average difference waves for the five artifact minimization approaches at Pz. (i) Scalp maps of the mean amplitude measured from 300–600 ms in the grand average difference wave. Error bars show the standard error of the mean. The VEOG-bipolar signals were computed as upper minus lower.

#### 3.3.1 Assessment of eyeblink confounds in the P3b data

We first assessed whether blinks differed across conditions and were therefore a potential confound. Figure 6b shows that the probability of blinks did not differ between targets and nontargets (t(39) = −0.52, p = 0.603, Cohen’s d’ = −0.083). Figure 6c shows the time course of the blink activity in the uncorrected VEOG-bipolar channel. The blink activity was earlier for nontargets than for targets, leading to a significantly more positive voltage for nontargets than for targets during the P3b measurement window (300–600 ms) (see Figure 6d; t(39) = 2.40, p = 0.021, Cohen’s d_z_ = 0.38). Thus, although eyeblinks occurred approximately equally often following targets and standards, the time course of the blinks differed, making them a potential confound.

Panels g, h, and i of Figure 6 show how blink activity distorted the target-minus-nontarget difference wave. When blinks were not corrected, a large negative deflection was present in the difference wave at FP1 and FP2 (panel g) beginning at approximately 300 ms. This is exactly what would be expected from the blink-related activity observed in the uncorrected VEOG-bipolar channel. The volume-conducted blink-related activity at FP1 and FP2 was dramatically reduced by all four approaches that included blink correction. Panel h shows the corresponding data from the P3b measurement channel (Pz). The greater blink-related activity for nontargets than for targets would be expected to partially cancel the typical P3b effect, and the difference wave was indeed smaller when no artifact correction was performed. The blink-related activity can also be seen as a negativity at anterior scalp sites in the scalp maps of the uncorrected data (panel i). Note that it is a negativity because the more positive blink-related voltage on nontarget trials was subtracted from the less positive blink-related voltage on target trials in the target-minus-nontarget difference wave.

Together, these results demonstrate that blinks are a potential confound in oddball tasks, which may alter the difference in amplitude between target and nontarget trials and distort the scalp distribution of the experimental effect. In the present data, blink-related activity reduced the amplitude of the target-minus-nontarget difference at Pz. If this difference in blink-related activity were smaller in one group or condition than in another, this could lead to an artifactual difference in P3b amplitude between these groups or conditions. In addition, there may be groups or conditions where blink-related activity is larger for targets than for nontargets, which would inflate the apparent magnitude of the P3b in the target-minus-nontarget difference wave.

#### 3.3.2. Effectiveness of ICA at minimizing eyeblink confounds in the P3b data

Figure 6e shows the grand average waveforms for the corrected VEOG-bipolar channel. There was only a 1.10 µV difference between the targets and nontargets during the P3b measurement window, which was not statistically significant (t(39) = −1.18, p = 0.244, Cohen’s d_z_ = −0.19). Moreover, given the ∼5% propagation of blinks to the Pz electrode (Lins et al., 1993), the confounding effect of eyeblinks on the P3b component at Pz would be less than 0.1 µV. Thus, the ICA-based correction was successful in minimizing the confounding EOG activity.

To provide additional information about the effectiveness of the blink correction procedure, we applied our semipartial correlation approach to the voltage during the P3b measurement window (300–600 ms) measured from the single-participant incorrect-minus-correct difference waves. We found that the semipartial correlation between the Pz channel and the corrected VEOG-bipolar channel was relatively small and not statistically significant (r(38) = 0.101, p = 0.542). This indicates that there was very little volume-conducted activity from the P3b component to the VEOG-bipolar channel. By contrast, the semipartial correlation between the uncorrected and corrected VEOG-bipolar signals was robust and statistically significant (r(38) = 0.583, p < 0.001). In other words, participants with larger differences in blink amplitude between targets and standards had larger differences in the corrected signal. In addition, the corrected VEOG-bipolar signal contained a noticeable deflection that approximately matched the time course of overall blink-related deflection in the uncorrected VEOG-bipolar signal. Together, these results indicate that ICA was not fully effective at minimizing blink-related activity. Nonetheless, it reduced the difference between targets and nontargets to nonsignificant levels that would be expected to produce a negligible effect when propagated to the Pz electrode site. Thus, ICA was sufficiently effective in the present dataset. However, it might not be sufficient in data with larger differences in blinking between target and nontarget trials.

#### 3.3.3 Assessment of data quality in the P3b data

Figure 7 shows the RMS(SME) values at the P3b measurement site (Pz) for each combination of the four scoring methods and the five artifact minimization approaches. As observed for the N400 component, ICA-based blink correction neither increased nor decreased the data quality. However, the three approaches that included rejection of trials with extreme values yielded better (smaller) RMS(SME) values. Thus, although artifact rejection reduced the number of trials (see Table 3), it still produced a net benefit in data quality. By contrast, the RMS(SME) for peak latency was largely unaffected by the four artifact minimization approaches (Figure 7d).

**Figure 7.**
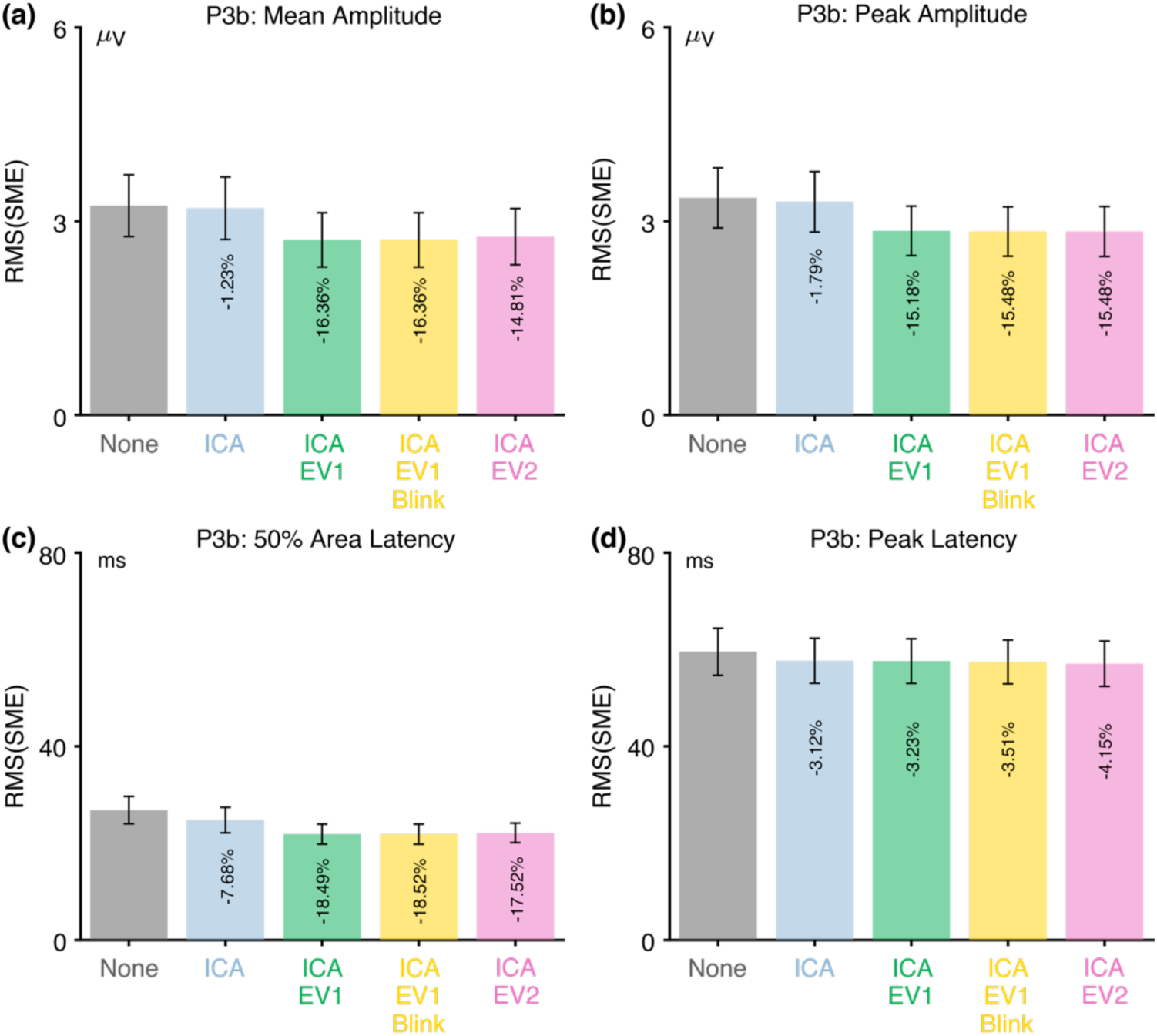
Root mean square of the standardized measurement error (RMS(SME)) from the P3b experiment for four different scoring methods and five different artifact minimization approaches. Smaller RMS(SME) values indicate higher data quality (less noise). Error bars show the standard error of the RMS(SME) values.

#### 3.3.4 Recommendations for the P3b

For studies like the ERP CORE P3b experiment, the present results demonstrate that blinks are a potential confound. We found that ICA-based blink correction reduced this confound to negligible levels, but we also found evidence that ICA did not fully eliminate all blink activity. Thus, researchers should carefully assess the effectiveness of ICA at removing blink confounds when it is applied to other datasets.

We also found that excluding trials with extreme values improved the data quality, with the reduction in number of trials being outweighed by the reduction in noise. There were no meaningful differences in data quality between the three rejection methods examined here. We have a slight preference for the ICA+EV1+Blinks approach, which is the most conservative and minimizes the possibility that participants were unable to see the stimulus on some trials because of blinks. These recommendations, along with a summary of the key P3b results, are provided in Table 4.

### 3.4 The mismatch negativity (MMN)

As shown in Figure 8a, a passive auditory oddball task was used to elicit the MMN. While participants viewed a silent video, they were exposed to a task-irrelevant sequence of standard tones (1000 Hz, 80 dB, p = .8, 800 trials) and deviant tones at a lower intensity (1000 Hz, 70 dB, p = .2, 200 trials). The MMN was measured from the deviant-minus-standard difference wave at the FCz electrode site.

**Figure 8.**
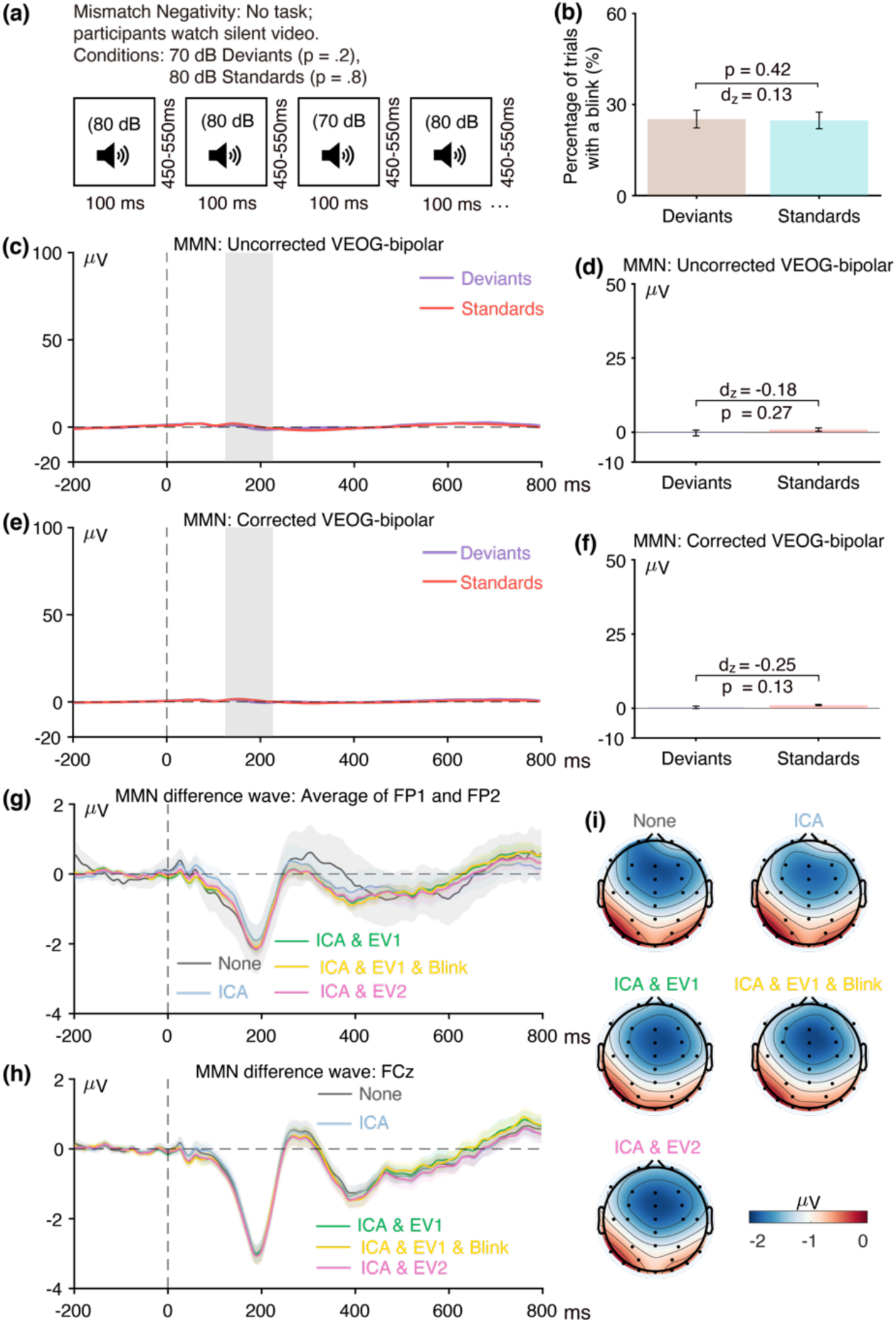
(a) Passive auditory oddball paradigm used to elicit the mismatch negativity (MMN). (b) Percentage of trials with a blink for the parent waves, measured from uncorrected VEOG-bipolar channel. (c) Grand average ERP waveforms for deviants and standards in the uncorrected VEOG-bipolar electrode site. (d) Mean amplitudes from the uncorrected VEOG-bipolar channel during the MMN measurement window for deviants and standards. (e) Grand average ERP waveforms for the deviants and standards in the corrected VEOG-bipolar channel. (f) Mean amplitudes from the corrected VEOG-bipolar channel during the MMN measurement window for the deviants and standards. (g) Grand average ERP difference waves (deviant minus standard) for the five artifact minimization approaches at the FP1 and FP2 electrode sites. (h) Grand average difference waves for the five artifact minimization approaches at FCz. (i) Scalp maps of the mean amplitude measured from 125–225 ms in the grand average difference wave. Error bars show the standard error of the mean. The VEOG-bipolar signals were computed as upper minus lower.

#### 3.4.1 Assessment of eyeblink confounds in the MMN data

Because the MMN stimuli were task-irrelevant, and the difference in intensity between deviants and standards was modest, we would not expect blinks to be triggered by the stimuli or blinking to differ between deviants and standard. Consistent with this expectation, Figure 8b shows that the probability of blinks did not differ between deviant versus standard tones (t(39) = 0.81, p = 0.42, Cohen’s d_z_ = 0.12). Similarly, Figures 8c and 8d show that there was no difference in the uncorrected VEOG-bipolar signal between deviant and standard tones (t(39) = - 1.12, p = 0.27, Cohen’s d_z_ = −0.18).

Figure 8e shows the grand average VEOG-bipolar channel after artifact correction, and Figure 8f shows the mean voltage in this channel during the MMN measurement window (125– 225 ms). The voltage was 0.783 µV more negative for standards than for deviants, but this difference was not statistically significant (t(39) = −1.55, p = 0.13, Cohen’s d_z_ = −0.25). As shown in panels g-i of Figure 8, the artifact minimization approaches that involved artifact correction had a relatively small impact on the deviant-minus-standard difference wave at the FP1 and FP2 electrodes and no discernible impact at the MMN measurement electrode (FCz).

These findings indicate that blinks were not a significant confounding factor in the ERP CORE MMN experiments.

#### 3.4.2. Effectiveness of ICA at minimizing eyeblink confounds in the MMN data

For the sake of completeness, we applied our semipartial correlation approach to the voltage during the MMN measurement window (125–225 ms) measured from the single-participant deviant-minus-standard difference waves. We found that the semipartial correlation between the uncorrected VEOG-bipolar channel and the corrected VEOG-bipolar channel was large and statistically significant (r(38) = 0.725, p < 0.001). This suggests that ICA was not fully effective at minimizing blink-related activity in the MMN paradigm. By contrast, the semipartial correlation between the FCz channel and the corrected VEOG-bipolar channel was relatively small and not statistically significant (r(38) = −0.027, p = 0.869). This suggests there was relatively little volume-conducted activity from the MMN component to the VEOG-bipolar channel.

Together, these results indicate that ICA was not completely effective at eliminating blink-related activity in the MMN paradigm, but that this was not problematic given that blinking did not differ between deviants and standards.

#### 3.4.3 Assessment of data quality in the MMN data

Figure 9 shows the RMS(SME) values at the MMN measurement site (FCz) for each combination of the four scoring methods and the five artifact minimization approaches. ICA alone had minimal impact on data quality, but the three approaches that included rejection of trials with extreme values produced a substantial decrease (>30%) in the RMS(SME) for the two amplitude scores. Three participants had extreme values on more than 25% of trials; excluding these participants improved the data quality even further for the amplitude scores (see supplementary Figure S11). The exclusion of trials with extreme values had minimal impact on the RMS(SME) for the two latency scores.

**Figure 9.**
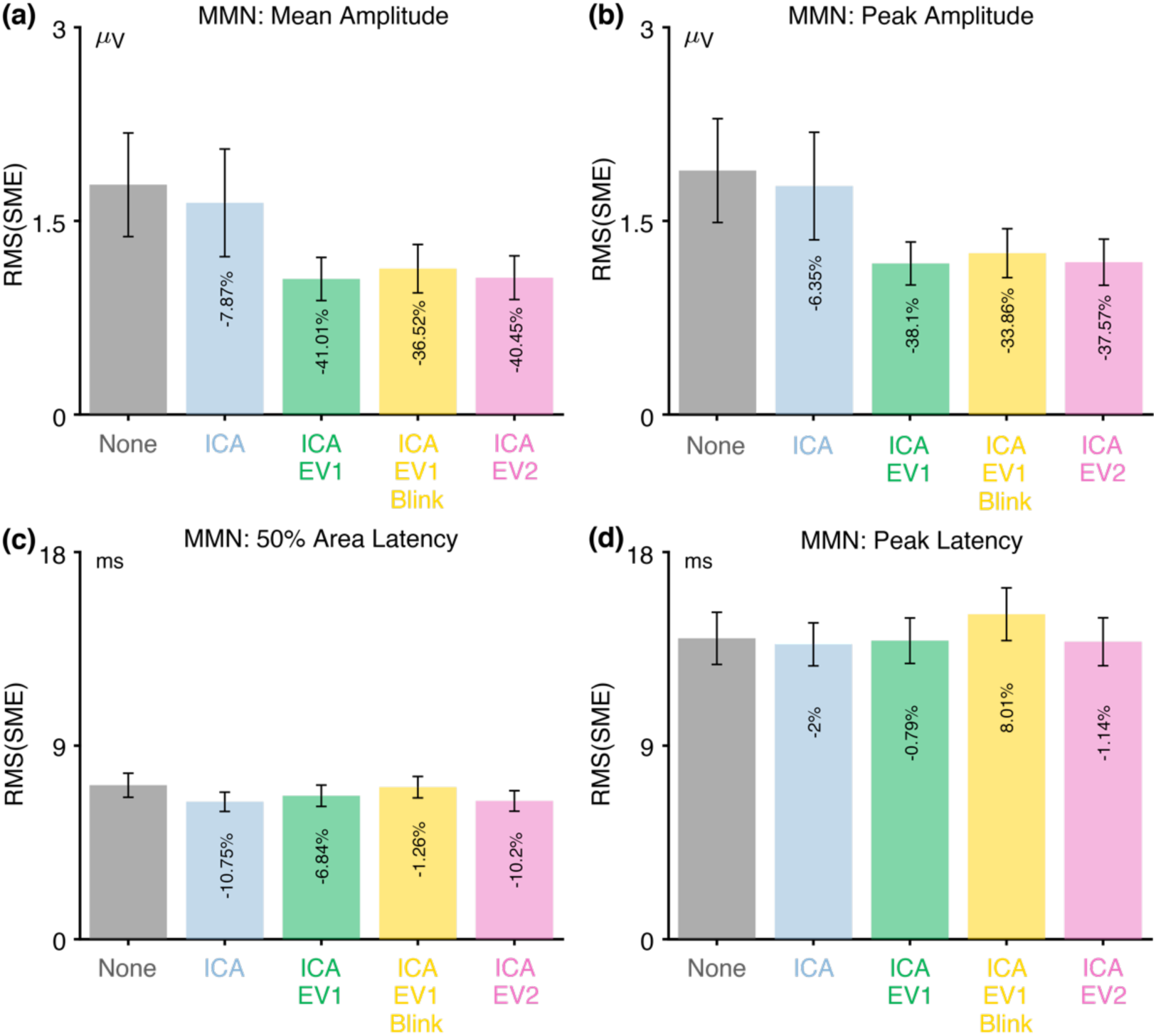
Root mean square of the standardized measurement error (RMS(SME)) from the MMN experiment for four different scoring methods and five different artifact minimization approaches. Smaller RMS(SME) values indicate higher data quality (less noise). Error bars show the standard error of the RMS(SME) values.

Whereas participants were instructed to withhold blinks until after they responded in the other ERP CORE paradigms, there were no behavioral responses in the MMN paradigm, so blinks could occur at any time in the MMN paradigm. As a result, the ICA+EV1+ blink approach led to the rejection of approximately 15% of trials (see Table 3), and the RMS(SME) values were elevated for this approach compared to the ICA+EV1 approach.

#### 3.4.4 Recommendations for the MMN

For studies like the ERP CORE MMN experiment, the present results indicate that blinks are unlikely to be a significant confound. ICA-based correction of blink artifacts is therefore not necessary, although it does not seem to have any negative consequences. Researchers may choose to include blink correction in their pipelines as a precautionary measure.

We recommend excluding trials with extreme values, which substantially improved the data quality in the present study. As with the previous components, we recommend the EV1 approach as the most conservative. However, we do not recommend excluding trials with blinks at the time of the stimulus. This is not necessary for the MMN paradigm given that the stimuli are auditory rather than visual, and exclusion of these trials degrades the overall data quality. These recommendations, along with a summary of the key MMN results, are provided in Table 4.

### 3.5 The N170 component

As illustrated in Figure 10a, the stimuli in the N170 paradigm consisted of a randomized sequence of faces, cars, scrambled faces, and scrambled cars. Participants were instructed to press one button for intact stimuli (faces or cars) and a different button for scrambled stimuli (scrambled faces or scrambled cars). There were 80 trials for each of the four stimulus classes. Here, we focus solely on the ERPs elicited by faces and cars. The N170 was measured from the faces-minus-cars difference wave at the PO8 electrode site.

**Figure 10.**
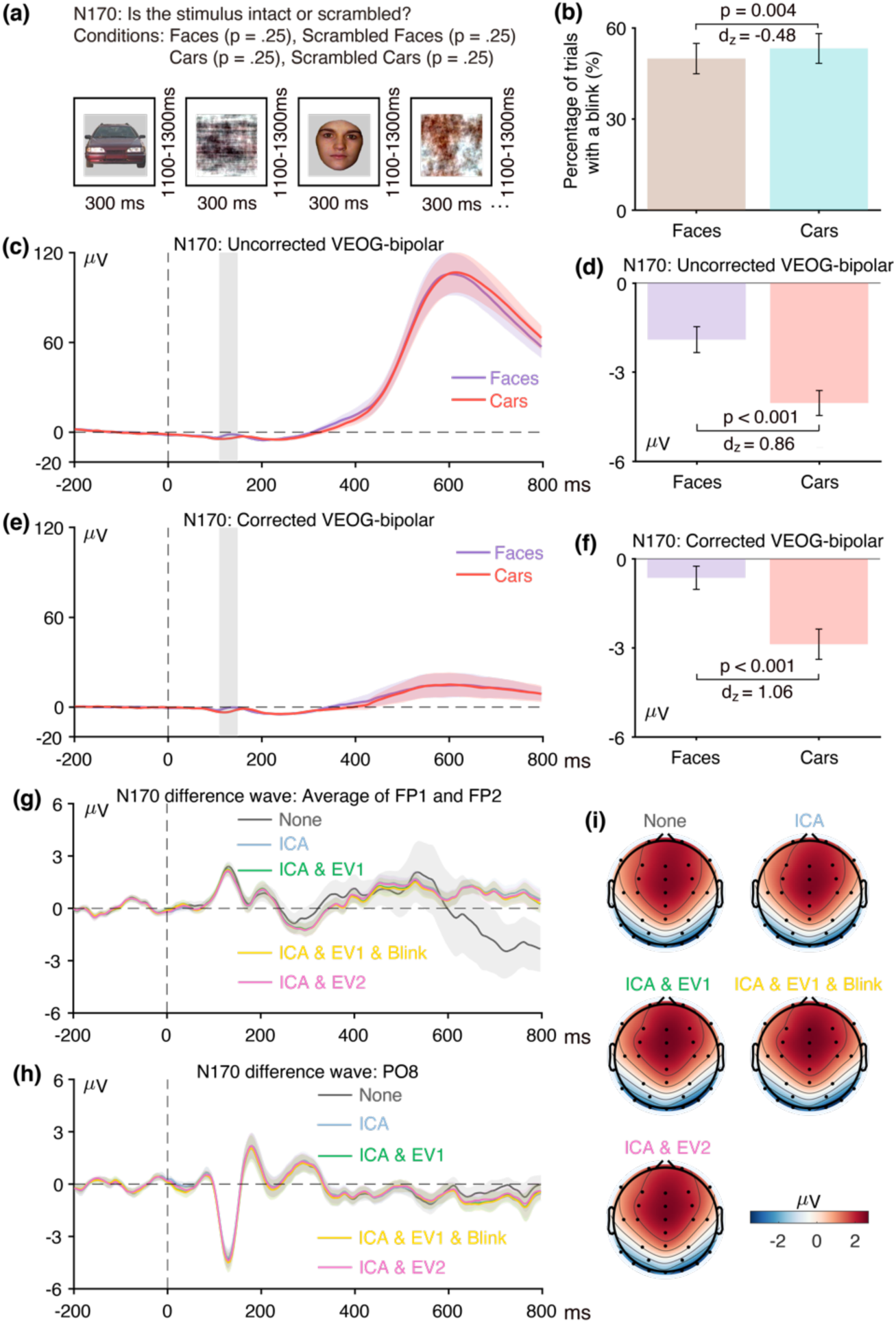
(a) Image categorization paradigm used to elicit the N170 component. (b) Percentage of trials with a blink for the parent waves, measured from uncorrected VEOG-bipolar channel. (b) Grand average ERP waveforms for faces and cars in the uncorrected VEOG-bipolar electrode site. (d) Mean amplitudes from the uncorrected VEOG-bipolar channel during the N170 measurement window for the face and car trials. (e) Grand average ERP waveforms for faces and cars in the corrected VEOG-bipolar channel. (f) Mean amplitudes from the corrected VEOG-bipolar channel during the N170 measurement window for the face and car trials. (g) Grand average ERP difference waves (face minus car) for the five artifact minimization approaches at the FP1 and FP2 electrode sites. (h) Grand average difference waves for the five artifact minimization approaches at PO8. (i) Scalp maps of the mean amplitude measured from 110–150 ms in the grand average difference wave. Error bars show the standard error of the mean. The VEOG-bipolar signals were computed as upper minus lower.

#### 3.5.1 Assessment of eyeblink confounds and the effectiveness of blink correction in the N170 data

Figure 10b shows that blinks were slightly but significantly more common for car trials than for face trials (t(39) = −3.06, p = 0.004, Cohen’s d_z_ = −0.484). Figure 10b shows the grand average waveforms for face and car trials in the uncorrected VEOG-bipolar channel. A very large deflection was present late in the epoch, presumably reflecting blinks that occurred after the behavioral response. This late activity was nearly identical on face trials and car trials. During the N170 measurement window (110–150 ms), however, the voltage was 2.13 µV more positive for faces than for cars, which was a significant difference (t(39) = 5.43, p < 0.001, Cohen’s d_z_ = 0.86). This difference was nearly the same size, 2.24 µV, in the corrected VEOG-bipolar signal and remained statistically significant (Figures 10e and 10f; t(39) = 6.72, p <0.001, Cohen’s d_z_ = 1.06). The unusual time course of this effect and the finding that it was not reduced at all by artifact correction suggests that it is a result of volume-conducted N170 activity rather than blink activity. Note that the N170 is positive at the front of the scalp because of the location and orientation of the N170 generator.

When we applied our semipartial correlation to the difference between face and car trials during the N170 measurement window, we found that the semipartial correlation between the PO8 channel and the corrected VEOG-bipolar channel was relatively robust and statistically significant (r(38) = −0.506, p = 0.001). This indicates that there was substantial volume-conducted N170 activity in the corrected VEOG-bipolar signal. Moreover, the semipartial correlation between the uncorrected VEOG-bipolar channel and the corrected VEOG-bipolar channel was small and not statistically significant (r(38) = 0.241, p = 0.139). These results provide additional evidence that the difference between faces and houses in the corrected VEOG-bipolar signal mainly reflects volume-conducted N170 activity rather than reflecting a blink confound.

However, the absence of a significant semipartial correlation between the uncorrected and corrected VEOG-bipolar signals is not definitive, so a close analysis of the time course of these signals is essential. Indeed, the large blink-related deflection that is visible late in the epoch in the uncorrected VEOG-bipolar signal (Figure 10c) is paralleled by a smaller deflection in the corrected VEOG-bipolar signal (Figure 10e), providing evidence that the correction was far from complete. However, the blink activity late in the epoch was reduced by over 80% in the corrected VEOG-bipolar signal relative to the uncorrected VEOG-bipolar signal, whereas the difference between faces and cars in the N170 latency range was no smaller in the corrected VEOG-bipolar signal than in the uncorrected VEOG-bipolar signal. This indicates that the effect during the N170 latency range primarily consists of volume-conducted N170 activity rather than residual EOG activity. Moreover, if any residual EOG activity remained, it would be reduced by over 96% after being propagated to the PO8 electrode site (Lins et al., 1993).

#### 3.5.2 Assessment of data quality in the N170 data

Figure 11 shows the RMS(SME) values at the a priori measurement site (PO8) for each combination of the four scoring methods and the five artifact minimization approaches. As was observed for the previous components, ICA neither increased nor decreased the RMS(SME) values. However, rejecting trials with extreme values improved the RMS(SME) values compared to the ICA-only approach and no artifact minimization (although the impact was negligible for the 50% area latency measure). Fewer than 4% of trials were excluded in any of the approaches that involved artifact rejection (see Table 3).

**Figure 11.**
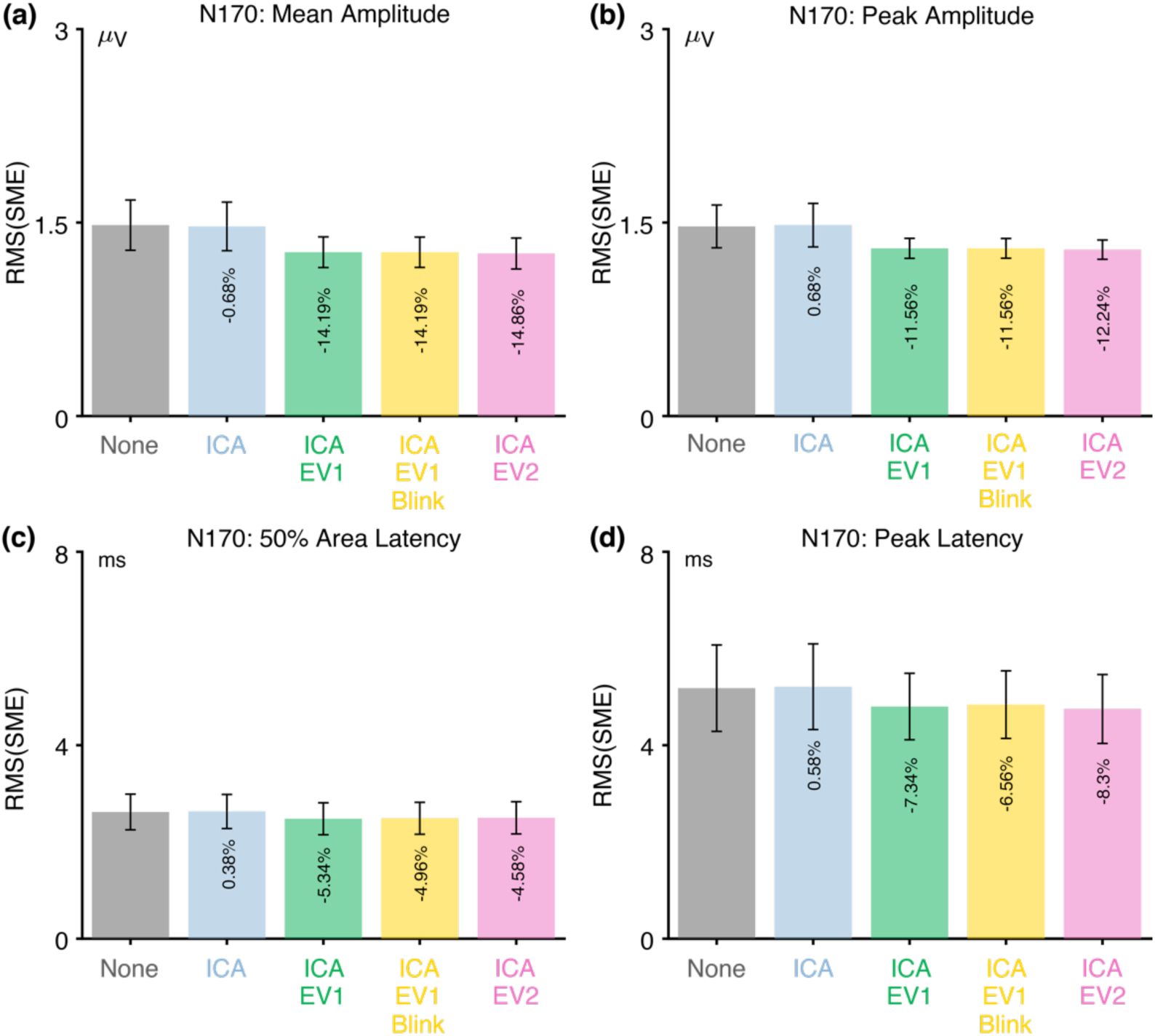
Root mean square of the standardized measurement error (RMS(SME)) from the N170 experiment for four different scoring methods and five different artifact minimization approaches. Smaller RMS(SME) values indicate higher data quality (less noise). Error bars show the standard error of the RMS(SME) values.

#### 3.5.3 Recommendations for the N170

For studies like the ERP CORE N170 experiment, the present results indicate that blinks are unlikely to be a significant confound, with any differences between conditions in the VEOG-bipolar signals being a result of volume-conducted N170 activity. ICA-based correction of blink artifacts is therefore not necessary, although it does not seem to have any negative consequences. Researchers may choose to include blink correction in their pipelines as a precautionary measure.

We recommend excluding trials with extreme values, which improved the data quality. There were no meaningful differences in data quality between the three rejection methods examined here. We have a slight preference for the ICA+EV1+Blinks approach, which is the most conservative and minimizes the possibility that participants were unable to see the stimulus on some trials because of blinks. These recommendations, along with a summary of the key N170 results, are provided in Table 4.

## 4. Discussion

In this study, we developed and applied a method for assessing the effectiveness of artifact minimization approaches in ERP research (summarized in Figure 1). This method focuses on two practical issues that arise from artifacts in many studies. First, blinks may differ across groups or conditions, producing a confound^9^, and it is important to ask whether any differences between groups or conditions in ERP measurements actually reflect residual EOG activity that was not completely removed by the artifact minimization procedure. This issue has been largely ignored by previous studies of the effectiveness of artifact correction methods. Second, artifacts may be a source of random, uncontrolled variance that reduce effect sizes and statistical power. Some previous methodology studies have examined how well artifact minimization procedures reduce this random noise using a variety of different approaches to quantifying the signal-to-noise ratio (Chang et al., 2020; Maddirala & Veluvolu, 2021; Phadikar et al., 2021; Yang et al., 2018). The present study used a new metric of data quality, the standardized measurement error, that is more directly related to effect sizes and statistical power (Luck et al., 2021). In general, we found that our artifact minimization approach was successful at both minimizing blink-related confounds and improving the signal-to-noise ratio across a broad set of ERP components.

### 4.1. Artifact minimization was effective in reducing confounds and improving data quality in the ERP CORE data

In the ERP CORE (Kappenman et al., 2021) data, we found that blinks are indeed a potential confound for several components, differing across experimental conditions in a way that could artificially inflate or decrease the observed experimental effects. We also demonstrated that a straightforward implementation of ICA-based artifact correction was reasonably successful at minimizing this confound (but was clearly not perfect). After correction, only the N400 component contained a statistically significant confound in the VEOG-bipolar channel that clearly reflected ocular activity, and this residual uncorrected EOG activity was small enough that it was likely to be negligible when propagated to the N400 measurement site (CPz). Thus, our relatively simple implementation of ICA-based blink correction was effective in solving the practical problem of blink-related confounds. To our knowledge, this issue has not been directly addressed in any previous studies of the effectiveness of ICA-based artifact correction.

It is important to note that we have focused on removing the electrooculographic artifact generated by the eyelid sliding over the cornea, but researchers may need to consider other potentials associated with blinks. For example, eyelid movements are produced by contraction of the orbicularis oculi muscles, which is accompanied by EMG activity. However, EMG phase is typically random, so this artifact is typically minimal in averaged ERPs. In addition, blinks are triggered by brain activity, and this brain activity would be difficult to remove via ICA.

We also found that combining artifact correction for blinks with artifact rejection for extreme voltages improved the data quality for four of the five ERP components, especially for amplitude scores. The improvement was considerable in some cases and modest in others. The only cases where this approach led to an apparent reduction in data quality were cases in which blink-related activity increased the size of the difference wave when artifact correction was not performed. In these cases, artifact correction is necessary to measure brain activity without distortion from ocular activity. Thus, the combination of artifact correction for blinks and artifact rejection for extreme voltages was valuable in almost every case, and it never produced a meaningful degradation in data quality. These findings provide evidence against the general proposal of Delorme (2023) that “EEG is better left alone,” at least in the context of the ERP CORE data.

We should note that one aspect of the present results was fully consistent with the findings of Delorme (2023). Specifically, we found that artifact correction had minimal impact on data quality in most cases. This suggests that blinks are a relatively small source of uncontrolled variance compared to other sources of noise. However, this may depend on the distance between the eyes and the measurement electrode, because blink-related voltages become progressively smaller with distance from the eyes (Lins et al., 1993). Indeed, the one case where we found that blink correction yielded a >10% improvement in data quality was the ERN, which was measured at a more anterior electrode site than most of the other components. Moreover, we found that blink correction was essential for minimizing blink-related confounds, an issue that was not considered by Delorme (2023). Thus, we would recommend that ERP researchers use blink correction in combination with rejection of trials with extreme values unless there is a compelling reason not to.

We have provided more specific recommendations for each of the five ERP components, which are summarized in Table 4. These recommendations should generalize to similar paradigms that are tested in similar participant populations using similar recording methods. Note that our claim is that the recommended approaches are reasonably effective and easy to implement, not that they are optimal. Future research can ask whether other approaches or parameters yield even better results.

### 4.2. Exclusion of participants with large numbers of artifacts

In the original ERP CORE analyses (and all studies of neurotypical young adults in our lab), participants were excluded from the final analyses if more than 25% of trials contained extreme values. This was relatively rare, leading to the exclusion of only one participant for most components (see supplementary Table S1). These participants were not excluded in the main analyses of the present study. However, supplementary analyses indicated that the aggregate group data quality was further improved when these participants were excluded (see supplementary Figures S7-S11).

Excluding participants also reduces the degrees of freedom for statistical analyses, and it is complicated to determine whether the increased data quality will offset the reduced degrees of freedom. In addition, the exclusion of participants has the potential to yield a nonrepresentative sample. Consequently, we are not making any specific recommendations about whether participants with large numbers of rejected trials should be excluded.

### 4.3. Assessing the effectiveness of artifact minimization with other datasets and procedures

It is unlikely that the artifact minimization procedures recommended here will be sufficiently effective for all experimental paradigms, all participant populations, and all recording methods. Indeed, we found that ICA-based blink correction often failed to completely remove blink artifacts. The residual artifactual activity was negligible in the present data but could be problematic in datasets that contain larger blink-related confounds. The results might also differ for studies in which participants were not instructed to withhold blinks until after making their behavioral response. Moreover, correction might not work as well in datasets with shorter recordings, higher noise levels, or different numbers of electrodes. In addition, the optimal approach to the rejection of trials with extreme values may differ across datasets. For example, higher rejection thresholds might be appropriate for infants and young children (Fló et al., 2022).

We have therefore tried to make it straightforward for other researchers to apply our assessment method to their own data. All elements of our method can be implemented in Matlab using a combination of EEGLAB, ERPLAB, and relatively simple scripts. In addition, we have made our scripts and data available at https://osf.io/vpb79/. We encourage other researchers to use this method to assess whether blinks (or other artifacts) are a potential confound in their data, how well their artifact minimization approach reduces any such confounds, and whether this approach actually helps their ultimate data quality (by reducing noise) or harms the ultimate data quality (by reducing the number of included trials). Researchers can also use our method to determine the optimal artifact minimization parameters for their data (e.g., the rejection thresholds that maximize data quality). Note, however, that we recommend using previous data to determine the optimal parameters and then applying these parameters in an a priori manner to new data.

The present method can also be used by methodologists to compare the effectiveness of different artifact minimization approaches. In particular, more advanced approaches to artifact correction such as artifact subspace reconstruction (Blum et al., 2019; Chang et al., 2020) and deep-learning based methods (Craik et al., 2019; Yang et al., 2018) will likely be more effective than the simple ICA implementation used in the present study. Note that our method could easily be adapted to other types of EEG-based analyses, such as time-frequency analysis and multivariate pattern analysis.

### 4.4. Different effects on amplitude and latency scores

The present results demonstrate that eyeblinks may differ across conditions in their timing and not just their amplitude. Consequently, they may be a confound for latency measures as well as amplitude measures, and some kind of eyeblink rejection or correction approach may be needed to eliminate this confound.

Although we found that the rejection of epochs with extreme values improved the data quality for amplitude scores in almost all cases, artifact rejection had little or no effect on data quality for latency scores. This suggests that it is more important to reject trials with extreme values when measuring amplitudes than when measuring latencies. Additional research is needed to determine why amplitude scores are more sensitive to epochs with extreme values and to determine whether some other kind of artifact rejection approach would be more valuable for latency scores.

More broadly, these results demonstrate that the value of a particular artifact minimization procedure may depend on how the ERP waveforms will ultimately be scored. Thus, there may be no single answer to the question of which artifact minimization approach is best. The optimal approach may depend on how the data will be scored, as well as the experimental paradigm, recording setup, and participant population. We have also found that the optimal filter settings vary across experimental paradigms and scoring methods (Zhang et al., 2023a, 2023b).

### 4.5. Imperfections of ICA-based artifact correction

Although we found that ICA-based artifact correction was sufficient to minimize confounds in the present data, we also found evidence that significant blink-related activity remained after correction for most of the components. Consequently, researchers cannot assume that this approach will be sufficient in very different datasets, especially those with larger differences between conditions in blink-related activity. We recommend that future studies examine differences between conditions and groups in the uncorrected and corrected VEOG-bipolar signals to determine whether blink-related confounds are present and, if so, whether they have been reduced to negligible levels by the correction process.

It should not be surprising that the present ICA-based blink correction approach was less than perfect. Indeed, it seems unlikely that any blink correction method will be perfect. The imperfection of the present approach likely stems from ICA’s assumption that the number of sources of activity is exactly equal to the number of data channels. This assumption is virtually always false (because it is extremely unlikely that the number of electrodes used in a given study will happen to exactly match the number of sources). When this assumption is violated, different true sources may be mixed together into a single independent component (IC), and a single true source may be split among multiple ICs (Groppe et al., 2009; Lindsen & Bhattacharya, 2010). Mixing and splitting of true sources may also occur if the blink activity does not exhibit zero-lag statistical independence from the EEG signals. If blink-related activity is not limited to a small number of ICs, and some blink-related activity remains in the ICs that are retained, not all of the blink-related activity will be removed in the correction process. This may explain why some evidence of residual blink-related activity remained after artifact correction.

The mixing and splitting of true components in ICA may also mean that the activity corresponding to an ERP component will be partially present in an IC that is being removed (Barbati et al., 2004; Castellanos & Makarov, 2006; Lindsen & Bhattacharya, 2010; Wallstrom et al., 2004). This could reduce differences between groups or conditions in this ERP component (see Ouyang et al., 2016 for evidence of this when ICA is used to remove the glosso-kinetic artifact). This is difficult to assess in real data, and our method does not directly address this possibility. Some studies using real EEG data combined with simulated eyeblink artifacts found that ICA did not remove any significant EEG activity when blinks were corrected (Frank & Frishkoff, 2007; Hoffmann & Falkenstein, 2008). However, we cannot be certain that this will be true in all datasets, independent of the nature of the data, the duration of the recording, and the number of channels. For this reason, we recommend a conservative approach in which only those ICs that clearly reflect artifacts are removed (e.g., an ICLabel probability of 0.8 or even higher).

### 4.6. Potential downsides of rejecting trials with extreme values

The rejection of trials with extreme values improved the data quality in the present study, with the benefits of reduced noise outweighing the cost of fewer trials. However, the rejection of trials may have a cost in terms of the interpretability of the data in some studies. When trials with extreme values or other artifacts are rejected, these trials essentially become missing data, and all the potential problems associated with missing data apply (Little & Rubin, 2019). In particular, when missing epochs are not randomly distributed across participants and trials, this could create bogus effects in some kinds of analyses (Heise et al., 2022). This issue is complex and beyond the scope of this paper, except that our method can be used to assess whether the proportion of rejected trials differs across groups or conditions. We should also note that fewer than 5% of trials were rejected because of extreme values in the present data, so any downsides of artifact rejection were likely minimal. However, a much higher percentage of trials might be rejected in other datasets, and researchers should think carefully about the potential for nonrandom rejection when larger numbers of trials are rejected.

## Supplementary Materials

### 1 Supplementary materials for all included subjects

#### 1.1. Artifact Minimization with Interpolation Instead of Rejection

When a channel briefly “goes bad,” it would be possible to interpolate the data for that channel rather than rejecting the trial. In theory, this would improve data quality by reducing the number of rejected trials. However, the interpolated data will not exactly match the true signal, so interpo-lation might reduce rather than increase data quality. In addition, if a trial is bad in one channel, it might also be bad in other channels, but perhaps without reaching the threshold for being marked as artifactual. This could result in interpolated values that far from the true values. In addition, if a channel is bad because of factors like movement artifacts, the data from the other channels on that trial might not be representative of a typical trial. Thus, it is an empirical question whether the value of keeping more trials outweighs the potential negative consequences of interpolation. We assessed this by testing two supplementary approaches that involved interpolation rather than rejection of channels with extreme values.

In the first supplementary approach, labeled “ICA & EV3”, we applied interpolation to an epoch when fewer than than 10% of the total number of channels were flagged for extreme values for that epoch. For these epochs, we used the data from all channels that were free from extreme values for that specific epoch to interpolate the flagged channels using the superfast spherical interpolation algorithm. An epoch was excluded from averaging when over 10% of the channels were flagged for extreme values.

The second supplementary approach, labeled “ICA & EV4” was identical except that we interpolated only the measurement channel for a given component (e.g., Pz for the P3b component) using the data of the artifact-free channels for that epoch. The epoch was excluded from averaging only if more than 10% of the channels (including the measurement channel) were flagged for extreme values. In other words, if more than 10% of channels were flagged, but the measurement channel was not flagged, the epoch was included during averaging. This was designed to exclude fewer trials.

**Figure S1:**
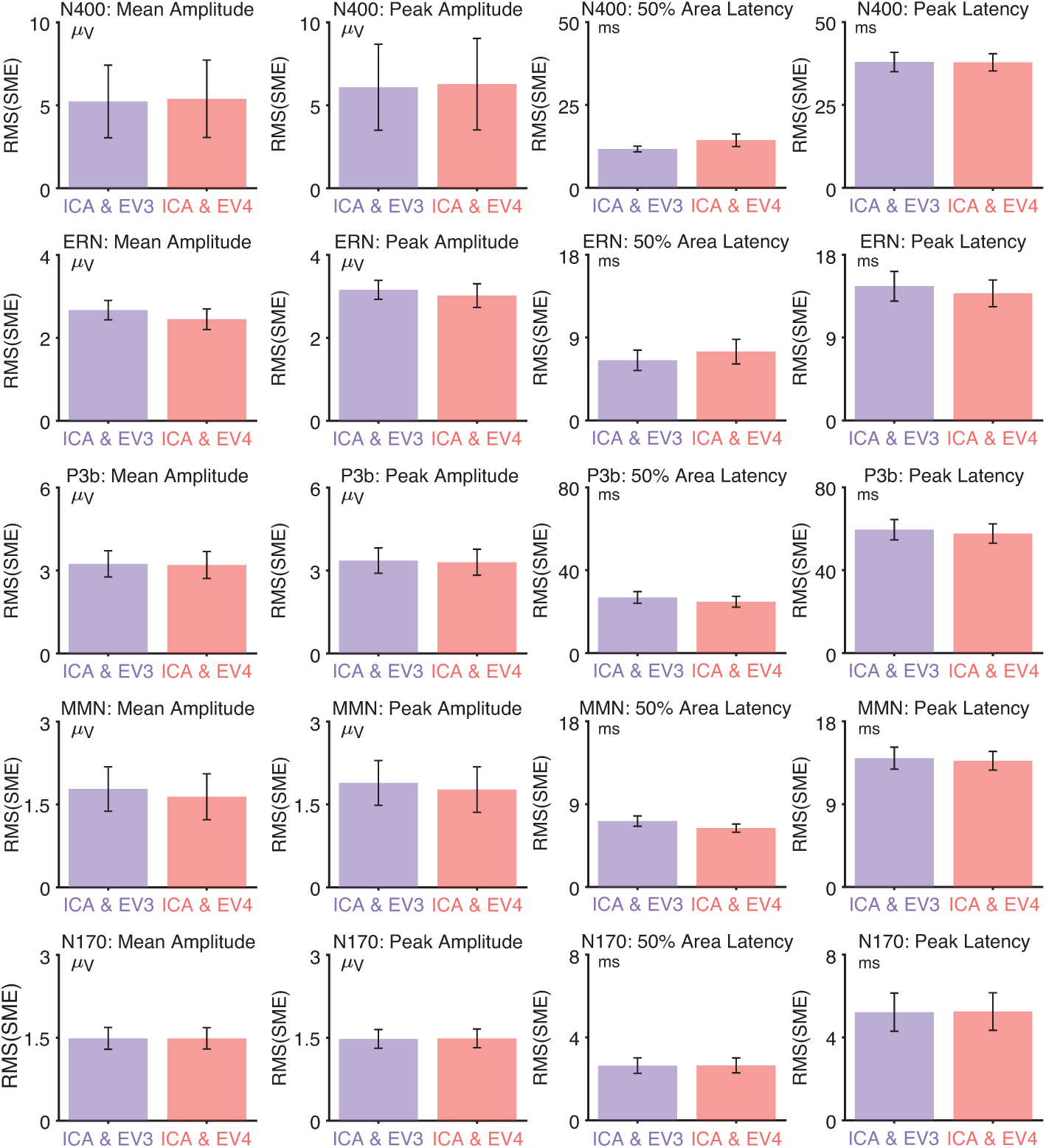
Root mean square of the standardized measurement error (RMS(SME)) for four di↵erent scoring methods and for five ERP components when using ICA & EV3 and ICA & EV4 artifact minimization approaches. Note that moving window peak-to-peak algorithm is with 100 µV and simple voltage threshold algorithm is with 200 µV. Error bars show the standard error of the mean.

#### 1.2. RMS(SME) for each combination of thresholds for each ERP component

**Figure S2:**
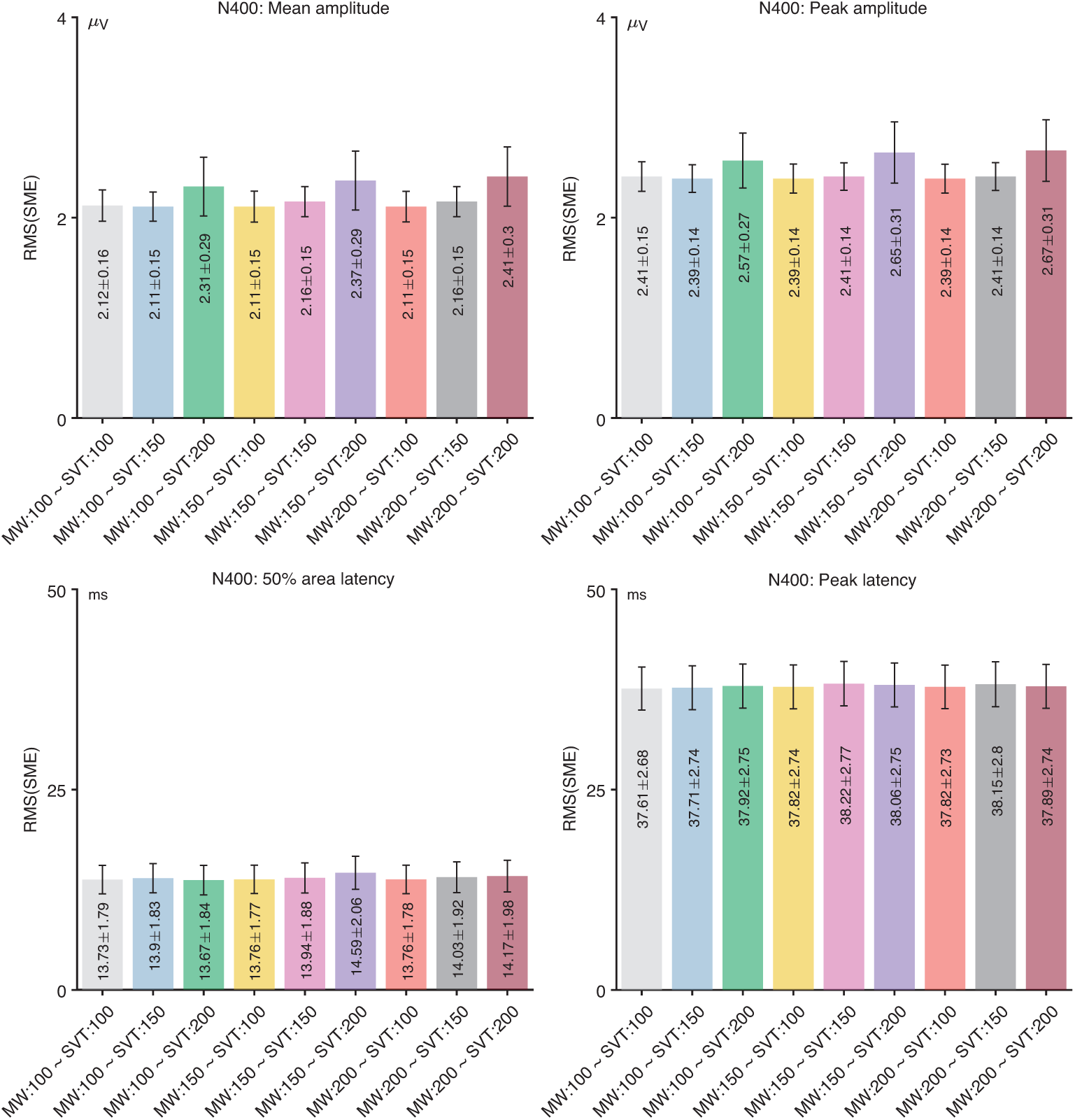
Root mean square of the standardized measurement error (RMS(SME)) from the N400 experiment for four different scoring methods and for each combination of thresholds for moving window peak-to-peak algorithm and simple voltage threshold algorithm. Error bars show the standard error of the RMS(SME) values.

**Figure S3:**
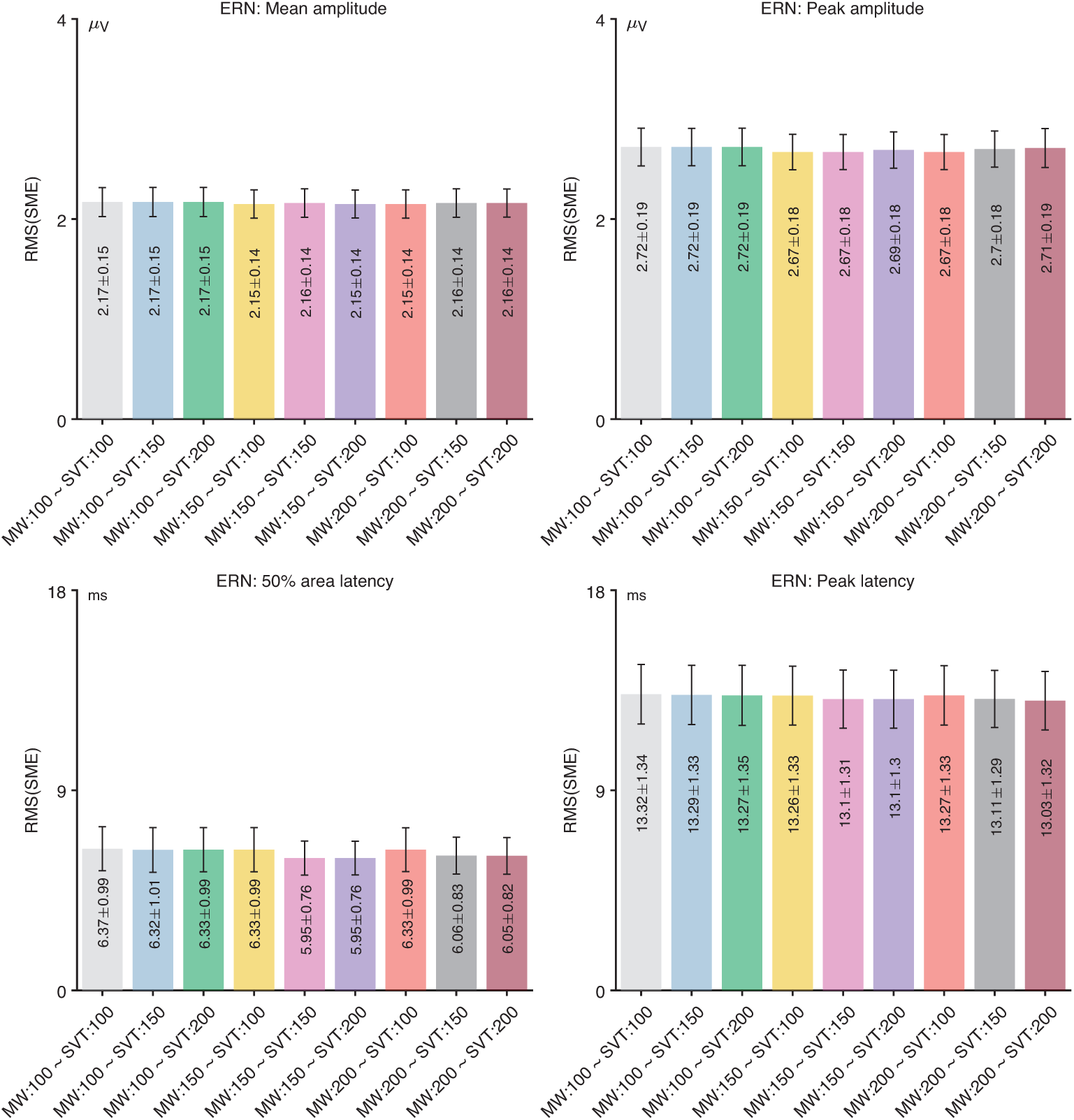
Root mean square of the standardized measurement error (RMS(SME)) from the ERN experiment for four different scoring methods and for each combination of thresholds for moving window peak-to-peak algorithm and simple voltage threshold algorithm. Error bars show the standard error of the RMS(SME) values.

**Figure S4:**
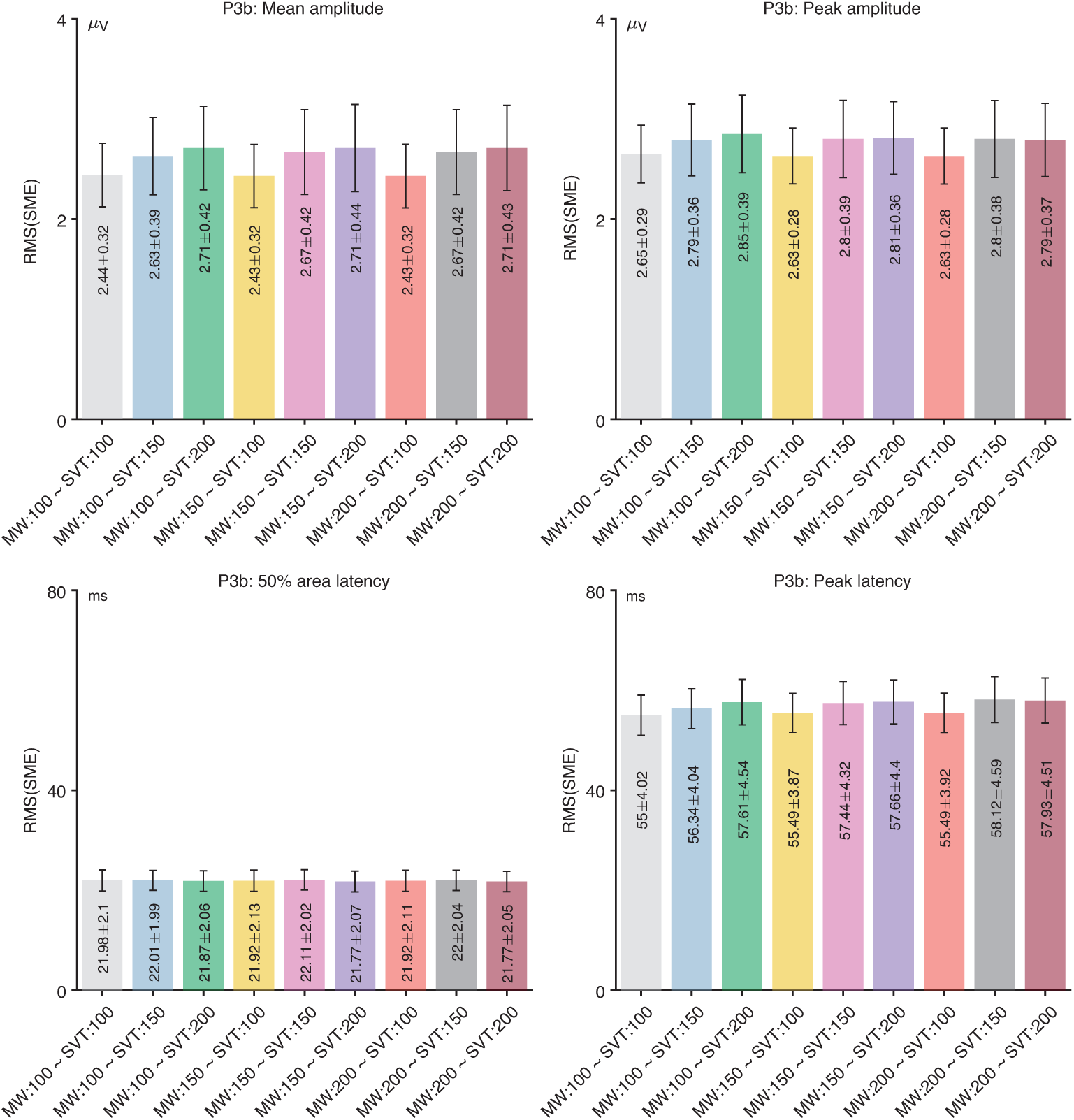
Root mean square of the standardized measurement error (RMS(SME)) from the P3b experiment for four different scoring methods and for each combination of thresholds for moving window peak-to-peak algorithm and simple voltage threshold algorithm. Error bars show the standard error of the RMS(SME) values.

**Figure S5:**
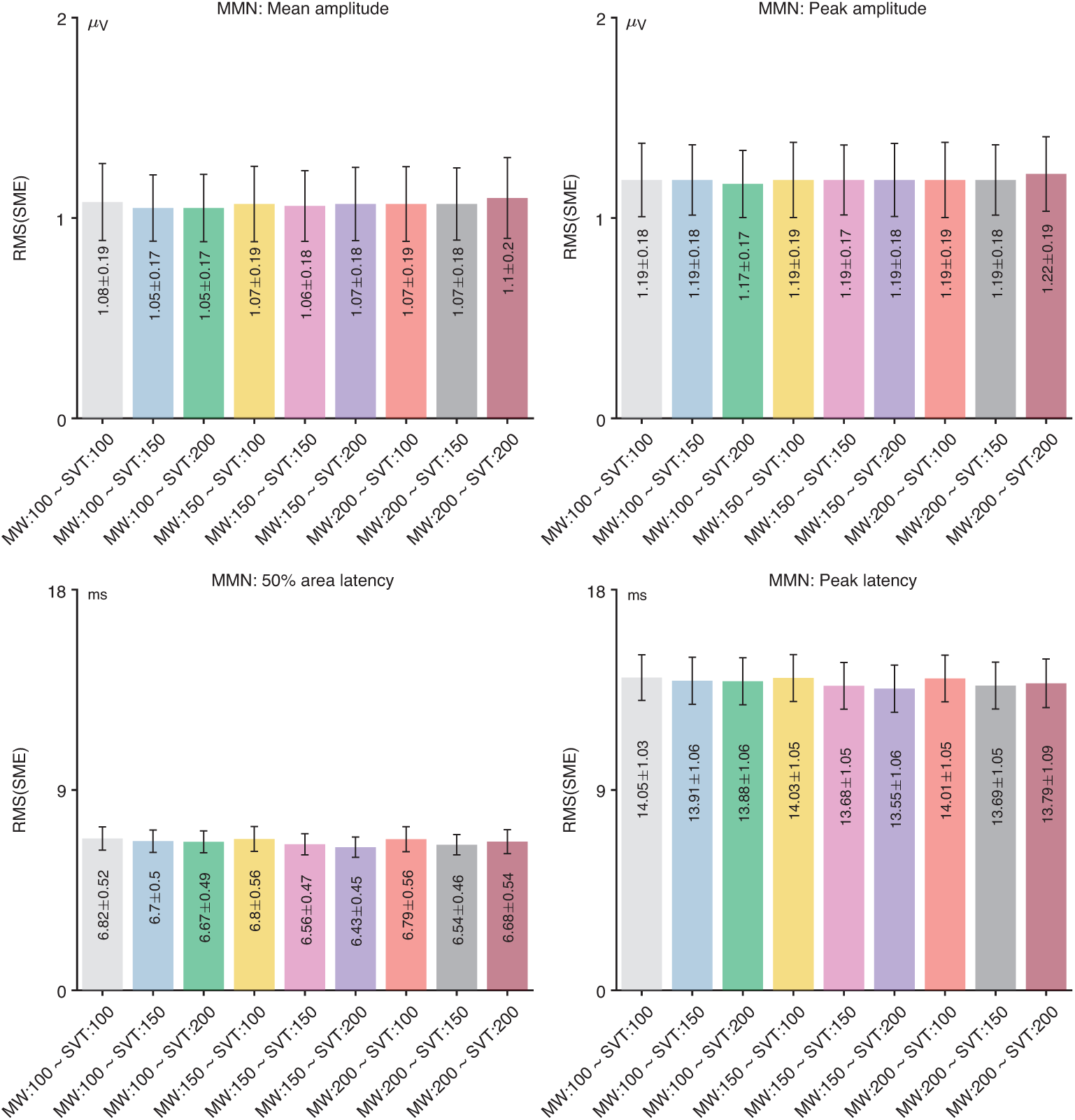
Root mean square of the standardized measurement error (RMS(SME)) from the MMN experiment for four different scoring methods and for each combination of thresholds for moving window peak-to-peak algorithm and simple voltage threshold algorithm. Error bars show the standard error of the RMS(SME) values.

**Figure S6:**
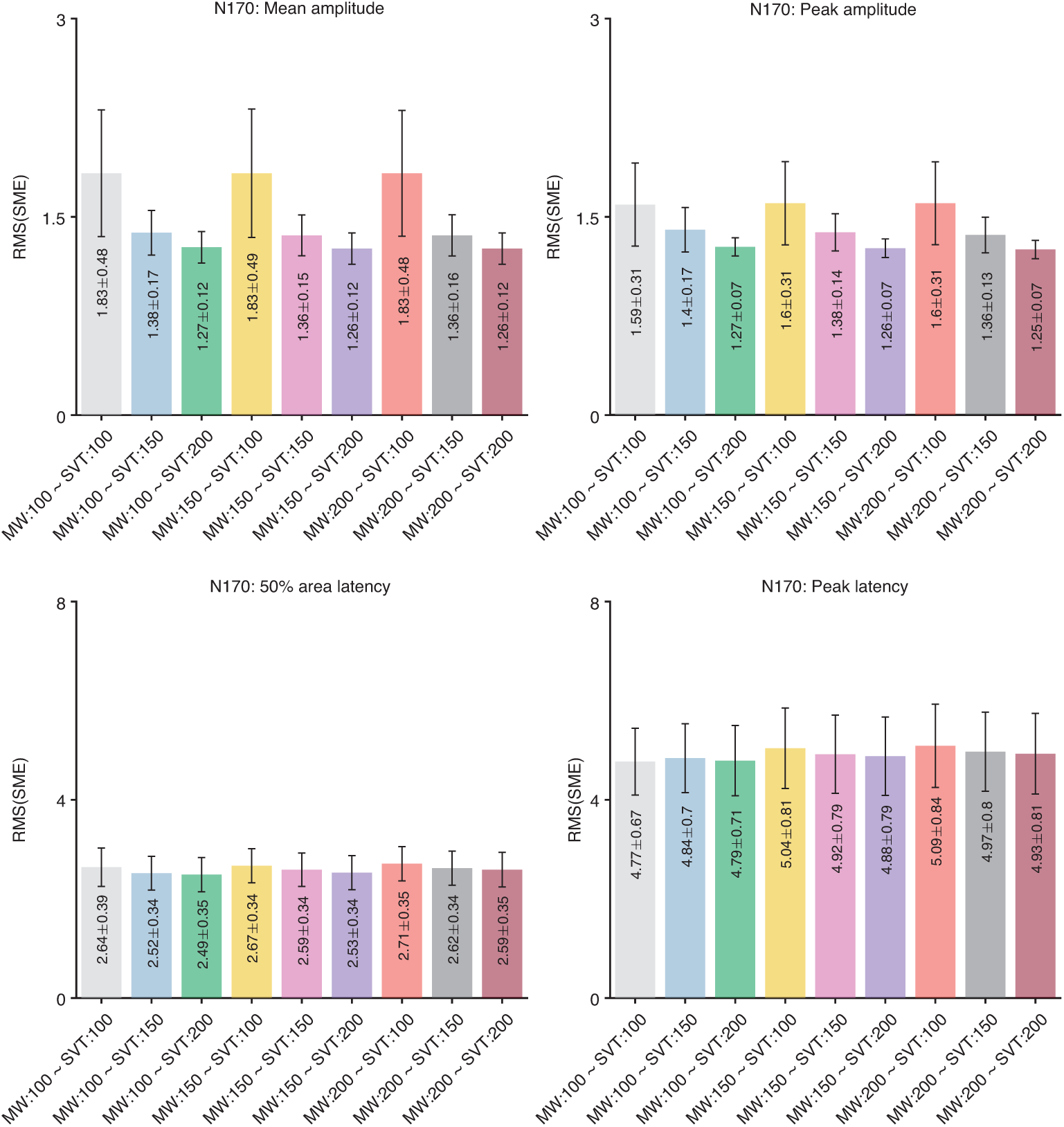
Root mean square of the standardized measurement error (RMS(SME)) from the N170 experiment for four different scoring methods and for each combination of thresholds for moving window peak-to-peak algorithm and simple voltage threshold algorithm. Error bars show the standard error of the RMS(SME) values.

### 2 Supplementary materials for exclusion of subjects with more than 25% rejected epochs

**Table S1:** Number of the excluded subjects that more than 25% of the total number of trials for both conditions were rejected. Note that One additional participant had two error trials was excluded from all ERN analyses.

|  | None | ICA | ICA & EV1 | ICA & EV1 & Blink | ICA & EV2 | ICA & EV3 | ICA & EV4 |
| --- | --- | --- | --- | --- | --- | --- | --- |
| *N400* | 0 | 0 | 1 | 1 | 0 | 0 | 0 |
| *ERN* | 0 | 0 | 1 | 1 | 0 | 0 | 0 |
| *P3b* | 0 | 0 | 1 | 1 | 0 | 0 | 0 |
| *MMN* | 0 | 0 | 3 | 6 | 1 | 1 | 1 |
| *N170* | 0 | 0 | 1 | 1 | 1 | 1 | 0 |

**Figure S7:**
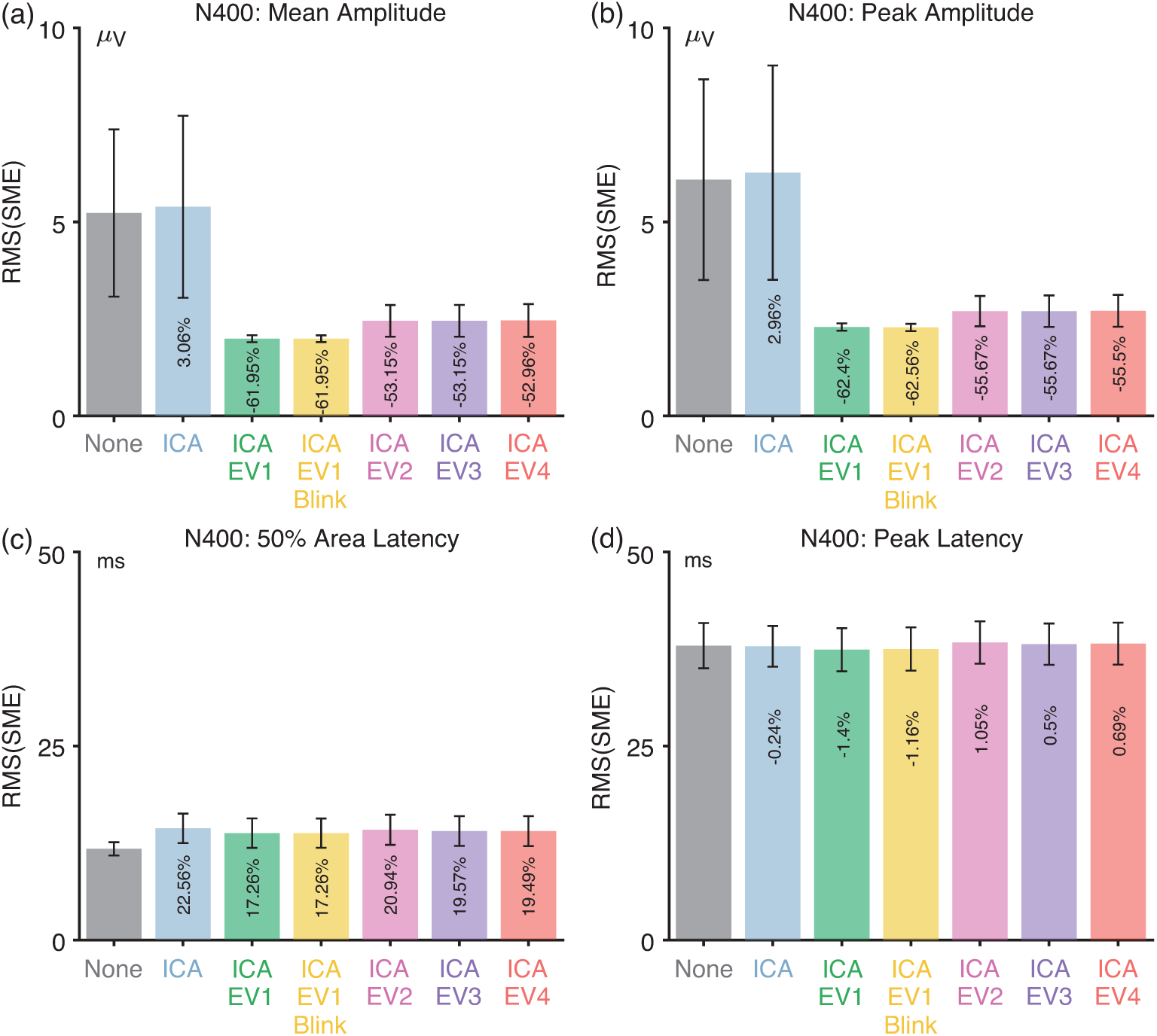
Root mean square of the standardized measurement error (RMS(SME)) from the N400 experiment for four different scoring methods and seven different artifact minimization approaches after excluding subjects with *>*25% rejected trials. Error bars show the standard error of the RMS(SME) values.

**Figure S8:**
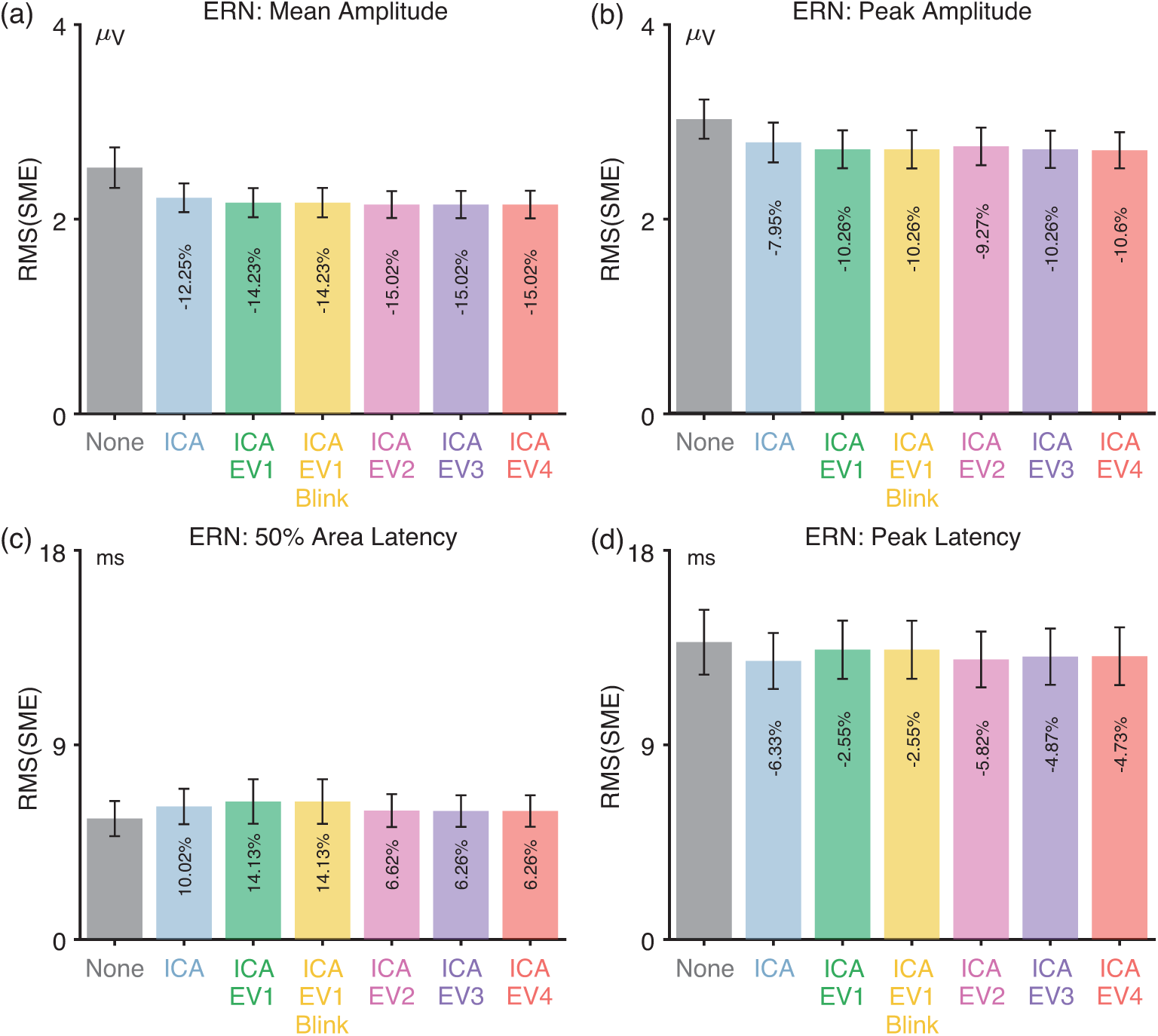
Root mean square of the standardized measurement error (RMS(SME)) from the ERN experiment for four different scoring methods and seven different artifact minimization approaches after excluding subjects with *>*25% rejected trials. Error bars show the standard error of the RMS(SME) values.

**Figure S9:**
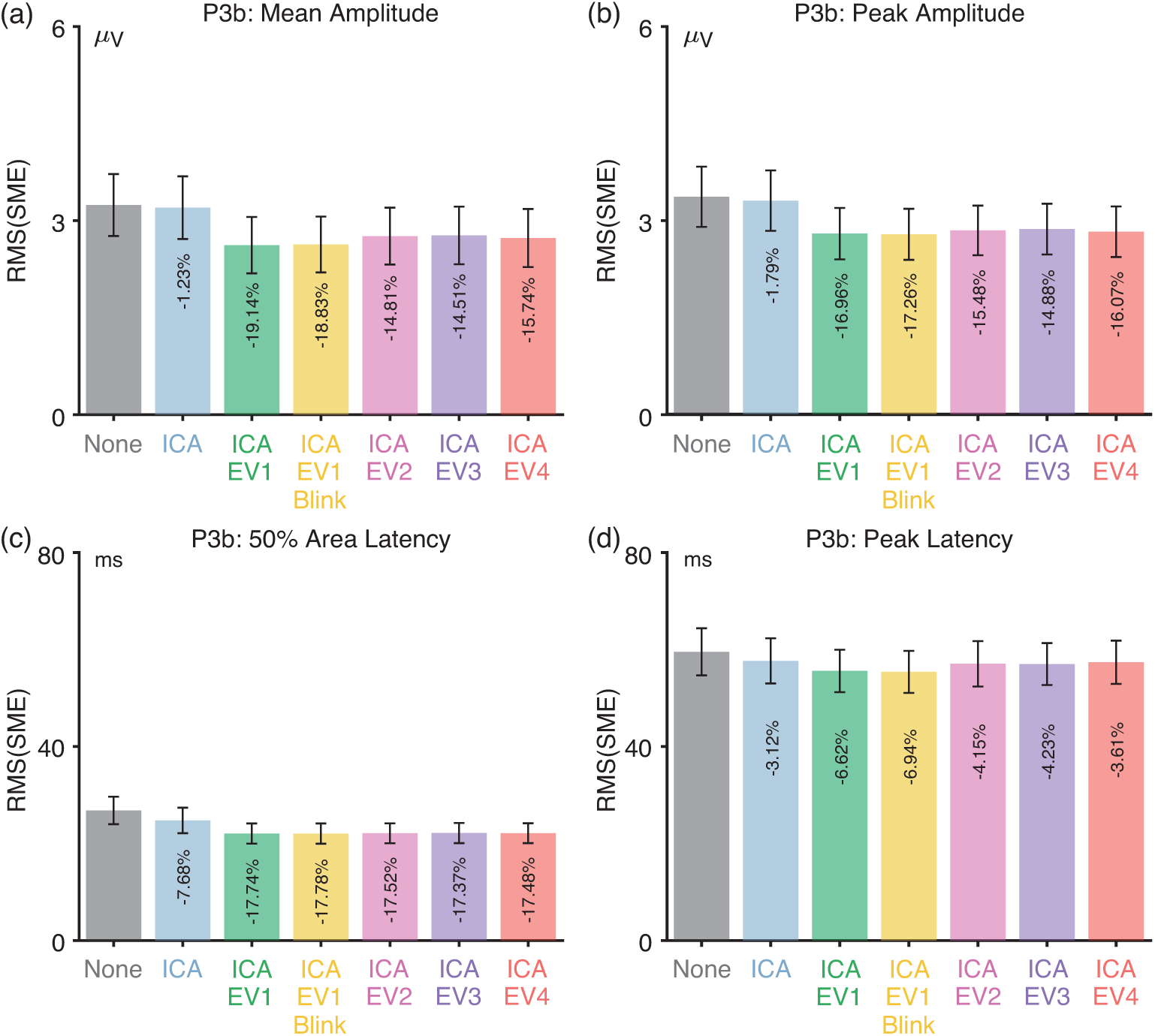
Root mean square of the standardized measurement error (RMS(SME)) from the P3b experiment for four different scoring methods and seven different artifact minimization approaches after excluding subjects with *>*25% rejected trials. Error bars show the standard error of the RMS(SME) values.

**Figure S10:**
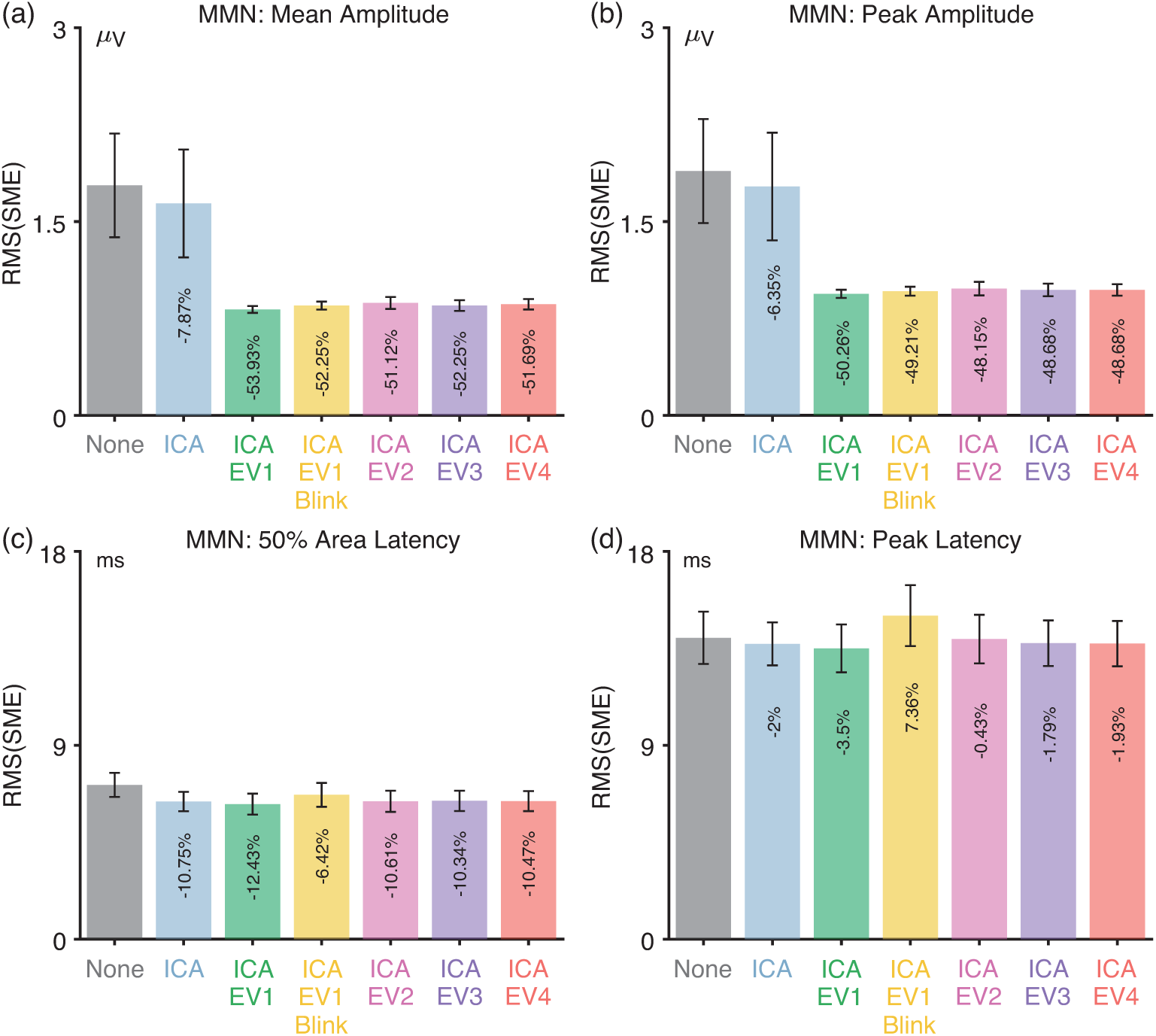
Root mean square of the standardized measurement error (RMS(SME)) from the MMN experiment for four different scoring methods and seven different artifact minimization approaches after excluding subjects with *>*25% rejected trials. Error bars show the standard error of the RMS(SME) values.

**Figure S11:**
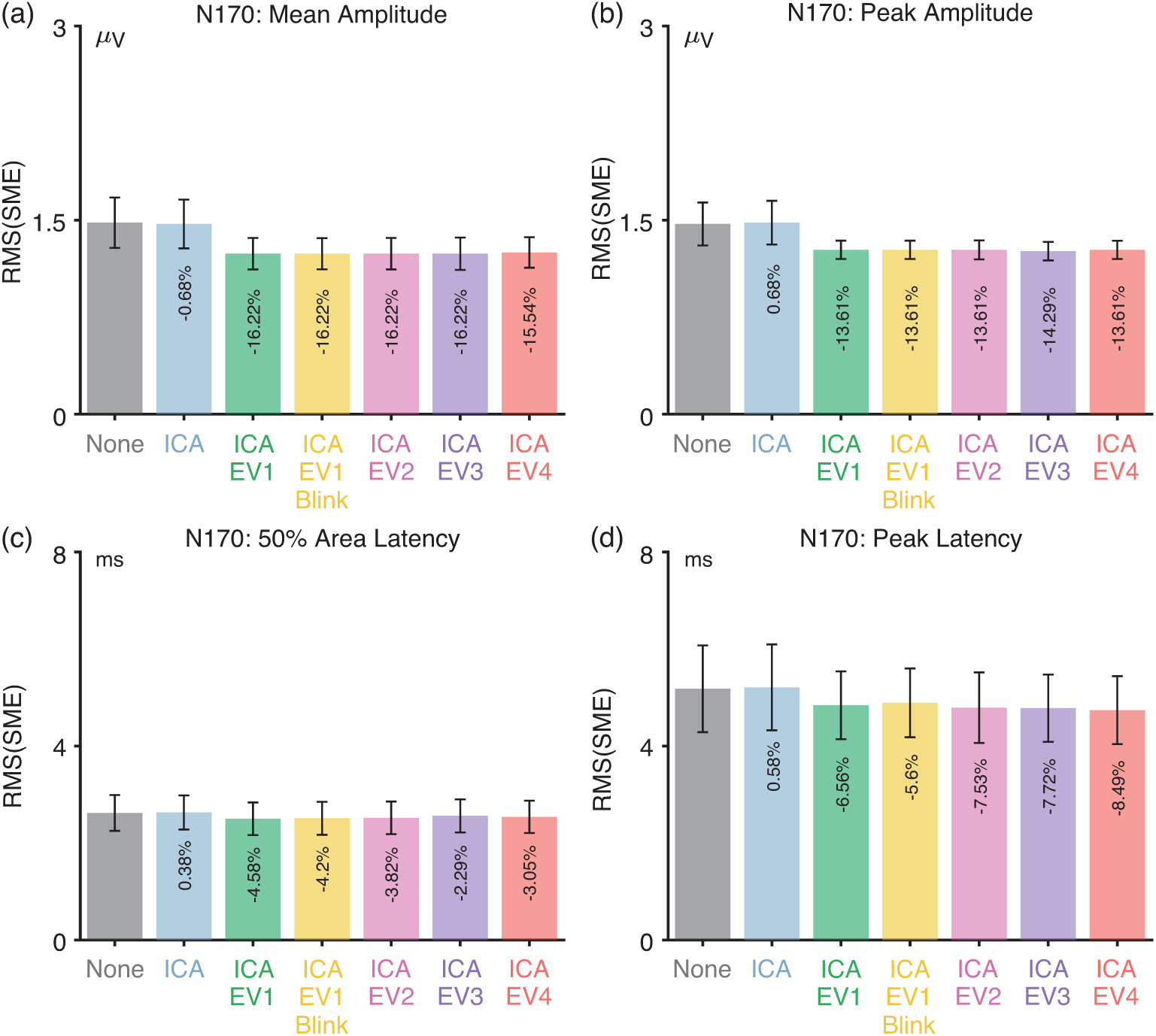
Root mean square of the standardized measurement error (RMS(SME)) from the N170 experiment for four different scoring methods and seven different artifact minimization approaches after excluding subjects with *>*25% rejected trials. Error bars show the standard error of the RMS(SME) values.

## Footnotes

1 N2pc and LRP are both isolated with contralateral-minus-ipsilateral difference waves, and this subtraction virtually eliminates eyeblink artifacts. Thus, blinks are a major issue for most components but not for N2pc and LRP. However, very small eye movements in the direction of the target stimulus (for N2pc) or response hand (for LRP) are a major concern for these components and require special analysis procedures that are not relevant for most other ERP components (Luck, 2014, 2022). We plan a separate paper assessing artifact minimization procedures for these components.

2 A rotation of the eyes during a blink may sometimes contribute to the artifact, but as reviewed by Lins et al. (1993), this is typically a minor contribution.

3 The bipolar VEOG is often computed with the reverse subtraction (lower minus upper). However, the upper-minus-lower subtraction is used here because it makes the polarity of any residual blink the same in the bipolar VEOG signal and the signals from the frontal and central scalp electrodes. In our data processing scripts, we initially computed the VEOG-bipolar signal as VEOG-lower minus FP2, and then we inverted this signal at the end of the processing sequence.

4 These ICs presumably picked up vertical eye movements as well, because the scalp distributions for blinks and vertical eye movements are quite similar. With 32 channels and very few vertical eye movements, we could not resolve separate ICs for blinks and vertical eye movements.

5 Hoffman and Falkenstein (2008) advocate for using mutual information rather than correlations, because mutual information allows for nonlinear relationships. We elected to use semipartial correlations primarily because we wanted an approach that would be easy for a broad range of researchers to implement. Moreover, the direction of a correlation (i.e., positive or negative) is useful in understanding the nature of the relationship. Finally, linearity is a reasonable approximation for the relationship between ocular artifacts and the signals recorded at scalp EEG electrodes.

6 Note that this is quite different from the regression-based blink correction approach of Gratton et al.(1983), which involves a separate regression for each participant using single-trial values. Our approach takes advantage of between-participant variance in the averaged waveforms, asking whether participants who have large voltages at CPz also have large voltages in the corrected VEOG-bipolar signal (after partialing out variance from the uncorrected VEOG-bipolar signal). This approach is used to assess the presence of uncorrected artifacts, not for performing artifact correction.

7 Another potential method for assessing the relationship between blinks and ERP activity is to record the bipolar electromyogram (EMG) from the orbicularis oculi muscle that is responsible for closing the eyelids (Marquardt et al., 2021). The amplitude of this signal provides a purer measure of blinks rather than a mixture of ERPs and blink-related voltages. Similarly, an eye tracker could be used to verify the timing of blinks. However, these measures were not recorded in the ERP CORE experiments.

8 This correlation did not cross the threshold for statistical significance (*p* = 0.0578) when a fixed ICLabel probability of >0.9 was used to classify ICs as blink-related, without manually inspecting ICs with lower probabilities.

9 We focused on blinks because they are the main artifacts that are likely to vary across conditions in the majority of ERP studies. Other kinds of artifacts may produce confounds in some experiments, and our method could be extended to these kinds of artifacts.

